# Spillover is the dominant non-photochemical quenching mechanism in angiosperms

**DOI:** 10.1101/2025.01.26.634902

**Authors:** Sanchali Nanda, Maximiliano Cainzos, Tatyana Shutova, Nazeer Fataftah, Verena Fleig, Jenna Lihavainen-Bag, Pushan Bag, Alfred R. Holzwarth, Stefan Jansson

## Abstract

Non-photochemical quenching (NPQ) is an important photoprotective process in plants, but all molecular details of the process(es) involved are yet not understood. We have used advanced spectroscopic techniques (including simultaneous time- and spectrally-resolved room-temperature chlorophyll fluorescence analysis and spectro-kinetic deconvolution) to analyse the processes in *Arabidopsis*, hybrid aspen and Scots pine plants. We used four well-characterized *Arabidopsis* lines (*npq1, npq2, npq4* and L17) affected in NPQ, together with hybrid aspen lines with corresponding modifications that we generated. The data are described best by a model for NPQ induction with up to five fluorescence components representing distinct biochemical entities. A dominant fluorescing species at the end of NPQ induction was identified as functionally detached and quenched LHCII but most importantly we believe that one represents a “PSII-PSI (Photosystem II-Photosystem I) complex” where direct energy transfer between PSII and PSI (spillover) take place. This provides strong quenching in all three plant species. We suggest a new integrated model for NPQ in higher plants where spillover is a major element and suggest roles for PsbS and zeaxanthin. Moreover, we discuss the link between NPQ and thylakoid rearrangements as thylakoid destacking facilitates direct contact between PSII and PSI; a prerequisite for spillover.

## Introduction

Non-photochemical quenching (NPQ) provides short-term adaptation to variations in factors such as light and temperature, as well as protection for the photosynthetic machinery of plants (Ruban, 2016). NPQ safely discards excess energy captured by photosynthetic pigments that cannot be used to drive photosynthesis. NPQ is essential to prevent damage to the photosynthesis apparatus which would cause losses in productivity or even death of tissues. However, if relaxation of NPQ is slow it may result in unnecessary dissipation of light, thereby compromising fitness and photosynthetic yields (Kromdijk *et al*., 2016).

NPQ involves several components and mechanisms, with some poorly understood details. The most rapidly induced component qE, which appears within seconds and relaxes within minutes, is linked to the PsbS protein (Li *et al*., 2000) and it acts on Light Harvesting Complex II (LHCII) (Holzwarth *et al*., 2009; Holzwarth and Jahns, 2014; Pawlak *et al*., 2020; Ruban and Wilson, 2021). A second important component, zeaxanthin (Zea), is formed in the light-driven xanthophyll cycle by violaxanthin epoxidase (VDE) (Demmig *et al*., 1987), and back-converted to violaxanthin by zeaxanthin epoxidase (ZE). It has been suggested that qE is modulated by Zea but a slower component, qZ, has also been linked to the presence of zeaxanthin (without involvement of PsbS) (Dall’Osto, Caffarri and Bassi, 2005; Nilkens *et al*., 2010). Very slowly relaxing components have also been described, including qH (whose action depends on plastid-located lipocalin (Malnoë *et al*., 2018)) and qI, photoinhibition caused by photosystem II (PSII) damage (Nawrocki *et al*., 2021).

Interest has also been regained in recent years in another mechanism – direct energy transfer between PSII and photosystem I (PSI) using Förster-resonance energy transfer - often named spillover. Spillover plays a prominent role in NPQ regulation in lichens (Slavov, Reus and Holzwarth, 2013), dinoflagellates (Slavov *et al*., 2016), red algae (Yokono, Murakami and Akimoto, 2011) and cyanobacteria (Akhtar *et al*., 2024). However, it is not reportedly present in diatoms (Flori *et al*., 2017). In vascular plants it has been argued that spillover is mechanistically unfavourable and unlikely because the very close contact between PSII and PSI units required for spillover conflicts with the traditional view of a strict lateral segregation of PSII-containing grana stacks and PSI-containing stroma-exposed thylakoid regions (Andersson and Anderson, 1980; Barber, 1980). However, thylakoid structure is not static and (re)organization is important for photosynthetic regulation (Johnson and Wientjes, 2020; Garty *et al*., 2024). We previously found that spillover is the main mechanism in the sustained quenching in conifer needles during the boreal winter and spring (Bag *et al*., 2020), and indications of a less prominent contribution of spillover to the regular light-induced NPQ. The presence of spillover has also been suggested – but not fully proven - in rice (Kim *et al*., 2023), spinach, cucumber, *Alocasia odora* (Terashima *et al*., 2024), implications of its presence have also been discussed (Yokono, Ueno and Akimoto, 2021). Spillover has not been studied in detail in *Arabidopsis*, but it has been suggested, for instance, that half of the PSII units in *Arabidopsis* might be in close contact with PSI (Yokono *et al*., 2015), which is consistent with a strong contribution to quenching.

Key players in NPQ have been identified using *Arabidopsis* mutants, including *npq1* (lacking VDE), *npq2* (lacking ZE), *npq4* (lacking PsbS), L17 (overexpressing PsbS) and mutants lacking one or more LHC proteins or PSII reaction centres (Johnson, 2020). Techniques commonly applied in NPQ characterization have included chlorophyll (Chl) fluorescence analysis methods, such as PAM (Pulse-Amplitude Modulated) (Johnson, Young and Horton, 1994; Li *et al*., 2004) and ultrafast time-resolved fluorescence analysis (Jahns and Holzwarth, 2012; Chukhutsina, Holzwarth and Croce, 2019; Yokono, Ueno and Akimoto, 2021).

We know a lot about NPQ regulation mechanisms, yet all details about sites of quenching, photophysical reactions, structures and kinetics of the processes involved are not clear. Complicating factors include the heterogeneity of the quenching and lack of adequate methods to separate the mechanisms and identify subprocesses involved. Each contributory process has specific onset and relaxation rates and may depend on numerous genetic and environmental variables, reviewed for example in (Ruban and Wilson, 2021).

Several reports suggest that short-term light adaptation involves structural reorganization of thylakoid membranes (Johnson and Wientjes, 2020; Rantala, Rantala and Aro, 2020; Ünnep *et al*., 2020). Light-induced thylakoid reorganization has also been characterized in the dinoflagellate *Symbodinium* in its “super-quenching state” (Slavov *et al*., 2016). On a longer time scale, in conifer needles thylakoid stacking is dramatically decreased during cold-induced sustained quenching, and lateral heterogeneity of the thylakoids almost disappears (Bag *et al*., 2020). Destacking in conifers has been linked to induction of strong spillover quenching as it requires thylakoids without pronounced grana-stacking, to enable the required physical interaction between PSII and PSI (Bag *et al*., 2020). It was believed that lateral heterogeneity of the thylakoids in higher plants evolved to prevent spillover by physically separating PSII and PSI units (Barber, 1980). However, recent literature has shown that the picture is more complex and also that strong thylakoid lateral heterogeneity architecture in higher plants is associated with dark adaptation, not necessarily representing normal light when NPQ takes place (Johnson and Wientjes, 2020; Garty *et al*., 2024). Hence, the relationship between NPQ and changes in thylakoid structure has attracted recent attention. However, it remains unclear which changes are causes and which are consequences of thylakoid reorganization.

Recently, new possibilities to study NPQ by analysing time- and wavelength-resolved (three-dimensional) quenching have been introduced (Nanda *et al*., 2024). This has several advantages over standard Chl fluorescence methods. For example, PAM analysis cannot generate defined spectra and concentrations of sub-components from total quenching signals. Time- and wavelength-resolution also enables specific component resolution with real-time recording of the quenching induction and relaxation processes, while ultrafast time-resolved techniques require long data accumulation times so only quasi-stationary (unquenched and quenched) states can be characterized (Holzwarth and Jahns, 2014). Three-dimensional *in vivo* chlorophyll fluorescence analysis of mutants lacking components of the NPQ machinery is expected to allow dissection of NPQ mechanism(s) and provide novel insights into its regulation. We here study a collection of well-characterized *Arabidopsis* mutants using time-dependent and spectrally resolved chlorophyll fluorescence (Nanda *et al*., 2024) and dissect the fluorescence quenching signal into various constituents thus characterizing the kinetic processes involved in the onset of quenching. By linking the data to electron microscopy studies, we also link them to thylakoid reorganization. We also generated a corresponding set of hybrid aspen (*Populus tremula x tremuloides* T89) lines and subjected them to the same quenching analysis to define general patterns or species differences and propose a new model for NPQ.

## Results

### Concentration profiles and assignment of spectral components reveal ***PSII*** and ***PSI*** *contributions to NPQ*

In a previous publication (Nanda et al. 2024) we used global target analysis (GTA) to dissect time-dependent and spectrally resolved room-temperature chlorophyll fluorescence spectra. Although GTA is a well-established method it comes with restrictions and therefore we opted for a more powerful and less restrictive method to deconvolute the spectra into its components; Multivariate Curve Resolution - Alternating Least Squares (MCR-ALS) analysis, see Material and methods for description and motivation. Spectra were in parallel subjected to GTA to complement/validate the results (Figs. S10-11) but only results from MCR-ALS are presented below.

The MCR-ALS spectro-kinetic analysis of *Arabidopsis*, hybrid aspen, and pine identified five Chl fluorescence components, represented by species-associated emission spectra (SAES), necessary to describe changes during NPQ induction (Figs. 1a, d, g). Their concentration profiles over time (Figs. 1c, f, and i) reveal the temporal dynamics of these components. The areas under the SAES, proportional to the relative fluorescence quantum yields, indicate their quenching contributions (Table 1).

**Fig. 1.**
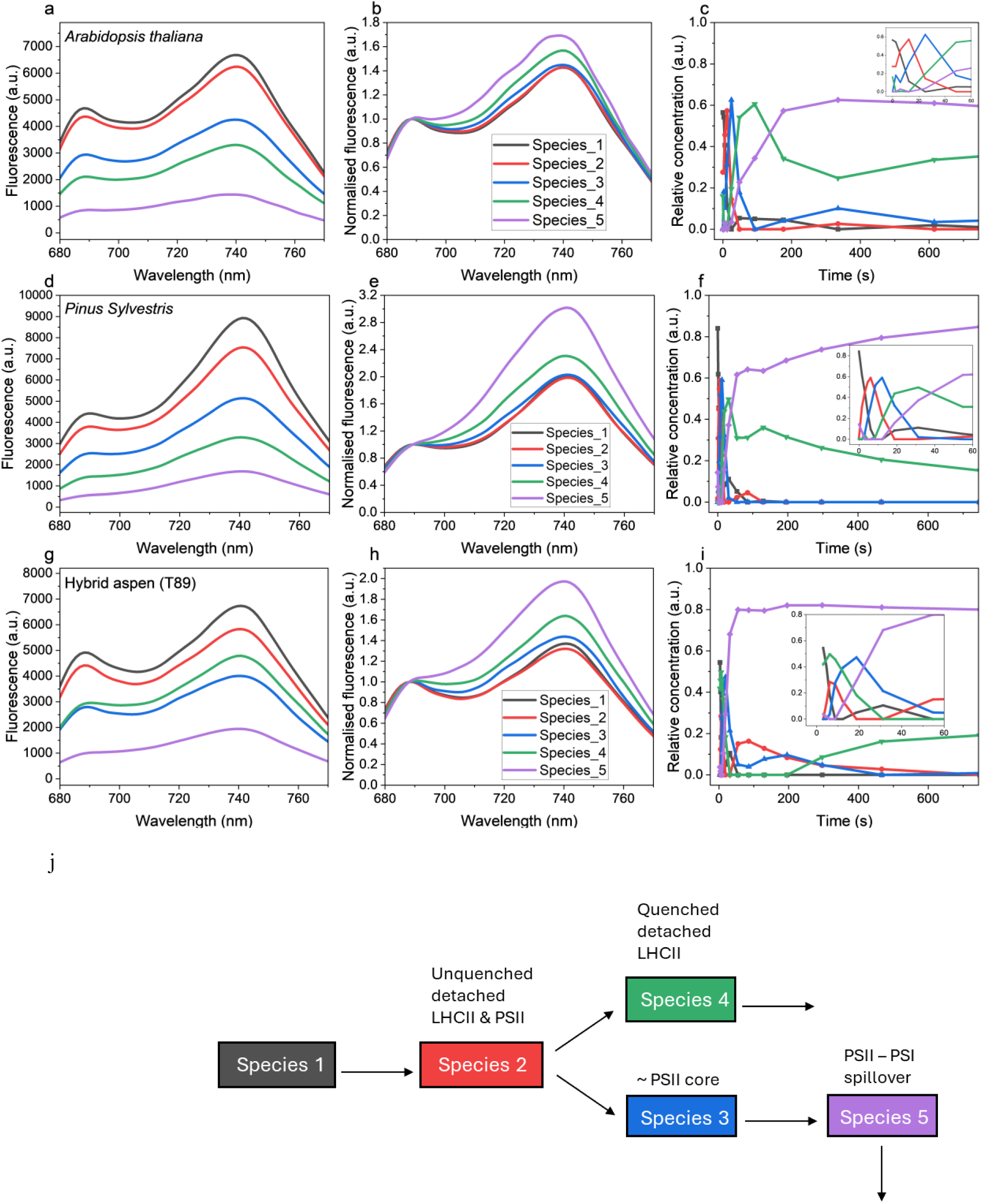
Results of decomposition of the 3-dimensional fluorescence spectra of the NPQ induction phase by MCR-ALS analysis into (**a**) SAES, (**b**) normalised SAES at 688nm and their (**c**) time-dependent concentrations for Col-0 (*A. thaliana*). The corresponding analysis for *P. sylvestris* in (**d**) – (**f**) and for hybrid aspen in (**g**) – (**i**). The NPQ for all samples was induced by 600 µmol m^−2^ s^−1^ red actinic light. Species 1 reflects the dominant (main) species in the dark-adapted state. The other species represent intermediate and end states. The resulting kinetic scheme as deduced from the concentration profiles is described in (**j**). The same colour code is used for representing the respective species in the various parts of the figure.

**Table 1.** Areas (relative to area of the most populated species 1 in the dark-adapted state) under the SAES of the different species. The area under the SAES is proportional to the fluorescence quantum yield of the respective species. Taken in relation to the starting species (dark-adapted state) the relative area gives the quenching factor (in parentheses) relative to the non-quenched species. *npq1\** & *vde\** are overnight dark-adapted samples. The error in the areas is ± 5%.

| Plant Species | Genotype | Relative SAES area (%) |  |  |  |  |
| --- | --- | --- | --- | --- | --- | --- |
|  |  | Species 1 | Species 2 | Species 3 | Species 4 | Species 5 |
| *Arabidopsis thaliana* | Col-0 | 100 | 94 (1.1) | 64.7 (1.5) | 48.9 (2) | 21.3 (4.7) |
|  | *npq4* | 100 | 86.7 (1.2) | 70.5 (1.4) |  |  |
|  | L17 | 100 | 77.5 (1.3) | 51.5 (1.9) | 33.6 (3) | 17.5 (5.7) |
|  | *npq1* | 100 | 85.8 (1.2) | 50.1 (2) | 55.3 (1.8) |  |
|  | *npq1* * | 100 | 84.9 (1.2) | 61.9 (1.6) | 53 (1.9) |  |
|  | *npq2* | 100 | 106.8 (0.9) | 76.1 (1.3) | 46.9 (2.1) | 18.2 (5.5) |
| T89 (hybrid aspen) | Wild type | 100 | 88.1 (1.1) | 60.1 (1.7) | 69.3 (1.4) | 27.2 (3.7) |
|  | *psbs* | 100 | 81.8 (1.2) | 88.2 (1.1) | 91.3 (1.1) |  |
|  | oePsbS | 100 | 94.9 (1.1) | 67.5 (1.5) | 43.8 (2.3) | 7.5 (13.3) |
|  | *vde* | 100 | 82.3 (1.2) | 102.1 (1) | 141.7 (0.7) |  |
|  | *vde* * | 100 | 85 (1.2) | 75.1 (1.3) | 50.3 (2) |  |
| *Pinus sylvestris* | Wild type | 100 | 85.8 (1.2) | 59.8 (1.7) | 37.9 (2.6) | 18.4 (5.4) |

The normalized spectra (Figs. 1b, e, h) showed that species 1–3 exhibited similar shapes, typical of PSII and LHCII fluorescence, with species 2 having the lowest contribution in the 700–740 nm range. In contrast, species 4 and 5 displayed distinct spectral features with pronounced far-red contributions. Species 4 did not have a pronounced contribution at 700 nm, but there were broad increases in its far-red contributions above 710-720 nm and peaks at both 688 and 740 nm, similar properties have been reported for quenched LHCII *in vivo* under NPQ conditions (Holzwarth *et al*., 2009; Miloslavina *et al*., 2011) or *in vitro* (Miloslavina *et al*., 2008; Ostroumov *et al*., 2020). For example, the quenched LHCII/PsbS complex co-reconstituted in proteoliposomes (Pawlak *et al*., 2020) has been suggested to have such properties. Therefore, we suggest that species 4 represent detached, and quenched, LHCII. Species 5 showed strongly increased contributions at 700 and 720 -740 nm, with a peak around 738-740 nm (and often a shoulder around 720 nm). These features are characteristic of PSI fluorescence (Tian *et al*., 2017; Bag *et al*., 2020). This increase in the far-red contribution of species 5 was even more pronounced in pine than in *Arabidopsis*. However, the species 5 band differed from a ‘classical PSI band’ as it also contained an intense band at 688 nm. The results from the analysis were similar for the three plants, and fluorescence species 5 was a highly quenched one with significant contributions from both PSII and PSI fluorescence, which we will refer to as the “PSII-PSI component” hereafter.

The temporal profiles (Figs. 1c, f, i) highlight the sequence of transitions during NPQ induction. At t = 0, the unquenched dark-adapted state was dominated by species 1, with minor contributions from species 2 and 4. Species 1 decayed rapidly within the first 15 seconds, coinciding with an increase in species 2, which peaked at ∼10 seconds. Subsequently, species 2 transitioned into species 3 and 4, which increased simultaneously. Species 3 reached its peak at ∼25, 15, and 20 seconds in *Arabidopsis*, pine, and aspen, respectively, before decaying as species 5 increased.

At the end of the NPQ induction phase, essentially only two strongly quenched species (4 and 5) were present, with species 5 accounting for ca. 80% of the total absorption cross-sections in pine and aspen and ca. 60% in *Arabidopsis* (Fig. 1c, f, i). Its spectral shapes in all three plants are similar and it is by far the most quenched species (judging from the area under the SAES and relative to the area of species 1’s SAES, cf. Table 1). The quenching factors of species 5 are 4.7, 3.7 and 5.4 in *Arabidopsis,* aspen and pine, respectively (Table 1). In *Arabidopsis* and pine, quenching gradually increased from species 1 (the unquenched dark-adapted state) to species 5, in aspen however species 3 was more quenched than species 4. At the end of the experiment, functionally detached but intermediately quenched LHCII provided ca. 40% of the total absorption cross-section in *Arabidopsis*, and somewhat less in pine and aspen.

In addition, species 4 oscillated slightly in *Arabidopsis* at around 100-200 s, and in pine at around 50-100 s but not at all in aspen. At longer times, the behaviour of species 4 and 5 differed somewhat between species. In *Arabidopsis*, species 5 slightly populated species 4 and as the concentrations of components 1-3 did not change significantly at this stage, species 4 was populated by some antennae splitting off from species 5. In pine, there was an opposite trend which we speculate comes from antenna movements which changes the absorption cross sections for different emitting Chl species.

The above results and interpretations suggest a sequence of reactions summarized in Fig. 1j. Species 1 transitioned into species 2, which then split into species 3 and 4. Species 3 subsequently decayed into species 5. While the behaviour of species 4 and 5 differed slightly across the three species, they consistently dominated the fluorescence profile at later stages in all three wild-type plant species. We need to point out that the biochemical assignment of species 2 is somewhat uncertain, presumably it is a mixture of several components (a pseudo-species). The scheme is very similar to the one we suggested based on analysis of early NPQ induction kinetics in *Arabidopsis* (Nanda *et al*., 2024) (using a similar measurement protocol and subsequent global target – not MCR-ALS – analysis).

### PsbS and zeaxanthin influence formation of the PSII-PSI spectral component

To see how major regulators influence NPQ reaction steps, intermediates, and end products we employed a set of well-characterized *Arabidopsis* mutants/transgenics and generated a corresponding set of hybrid aspen lines. A basic characterization of the aspen lines is presented in Fig. S4. Various lines were tested to check their PsbS protein expression levels. The lines with the highest and lowest PsbS expression are represented in Figs. S4a-b and were chosen for further analysis. These lines and the aspen *vde* mutants had similar NPQ induction and relaxation behaviour to the corresponding *Arabidopsis* mutants (Fig. S4d). None of the aspen lines had any obvious deviations in growth phenotype from wild type (WT) (Fig. S5).

*Arabidopsis npq4*, lacking PsbS, developed little NPQ (values of ca. 0.5-0.6, Fig. S6) and displayed slow biphasic induction kinetics and relaxation (as previously reported (Li *et al*., 2000)). The corresponding aspen mutant behaved similarly, with even lower NPQ (Fig. S8). According to the MCR-ALS analysis, three and four kinetic components or species were sufficient to describe well the observed kinetic development of *Arabidopsis* and aspen mutant spectra, respectively. The corresponding SAES had similar shapes and showed limited (ca. 30%) quenching, consistent with these mutants’ lower overall quenching. Plants lacking PsbS had no fast (≤10-15 s halftime) kinetic components giving rise to detached quenched LHCII, consistent with earlier data on the absence of functionally detached and quenched LHCII in *npq4* by ultrafast time-resolved fluorescence experiments (Holzwarth *et al*., 2009). The fastest processes (formation of species 3) in both plant species, with half-times of ca. 50 s in aspen and 100 s in *Arabidopsis*, resulted in only slightly quenched species with no pronounced far-red increase in the SAES (Figs. 2 and 3). Slower (over 500 s) processes were associated with minor SAES changes and very little quenching. In both species, lack of PsbS resulted in the development of SAES very similar to the dark state SAES, reflecting unquenched PSII supercomplexes. Thus, PsbS was necessary for the formation of two entities: detached quenched LHCII and the PSII-PSI component of NPQ. Accordingly, the NPQ spectra of PsbS mutants were more or less flat (Fig. S9), except for a small peak at 680 nm in the *Arabidopsis npq4* spectrum (Fig. S7)

**Fig. 2.**
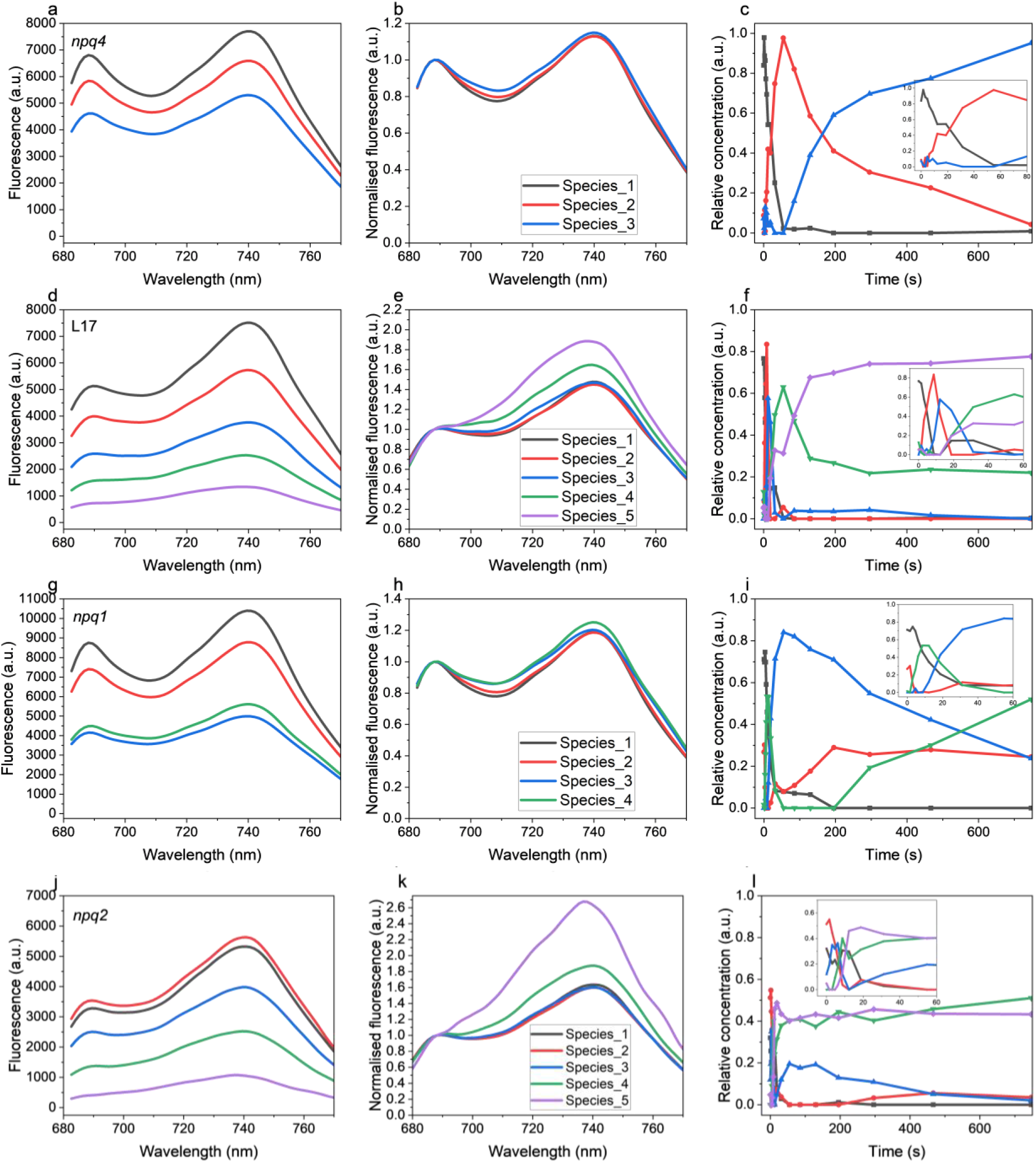
MCR-ALS analysis for *Arabidopsis* genotypes as SAES, normalised SAES at 688nm and time dependent concentration profiles respectively for *npq4* (**a**, **b**, **c**); L17 (**d**, **e**, **f**); *npq1* (**g**, **h**, **i**) and *npq2* (**j**, **k**, **l**). NPQ for all samples was induced by 600 µmol m^−2^ s^−1^ red actinic light.

**Fig. 3.**
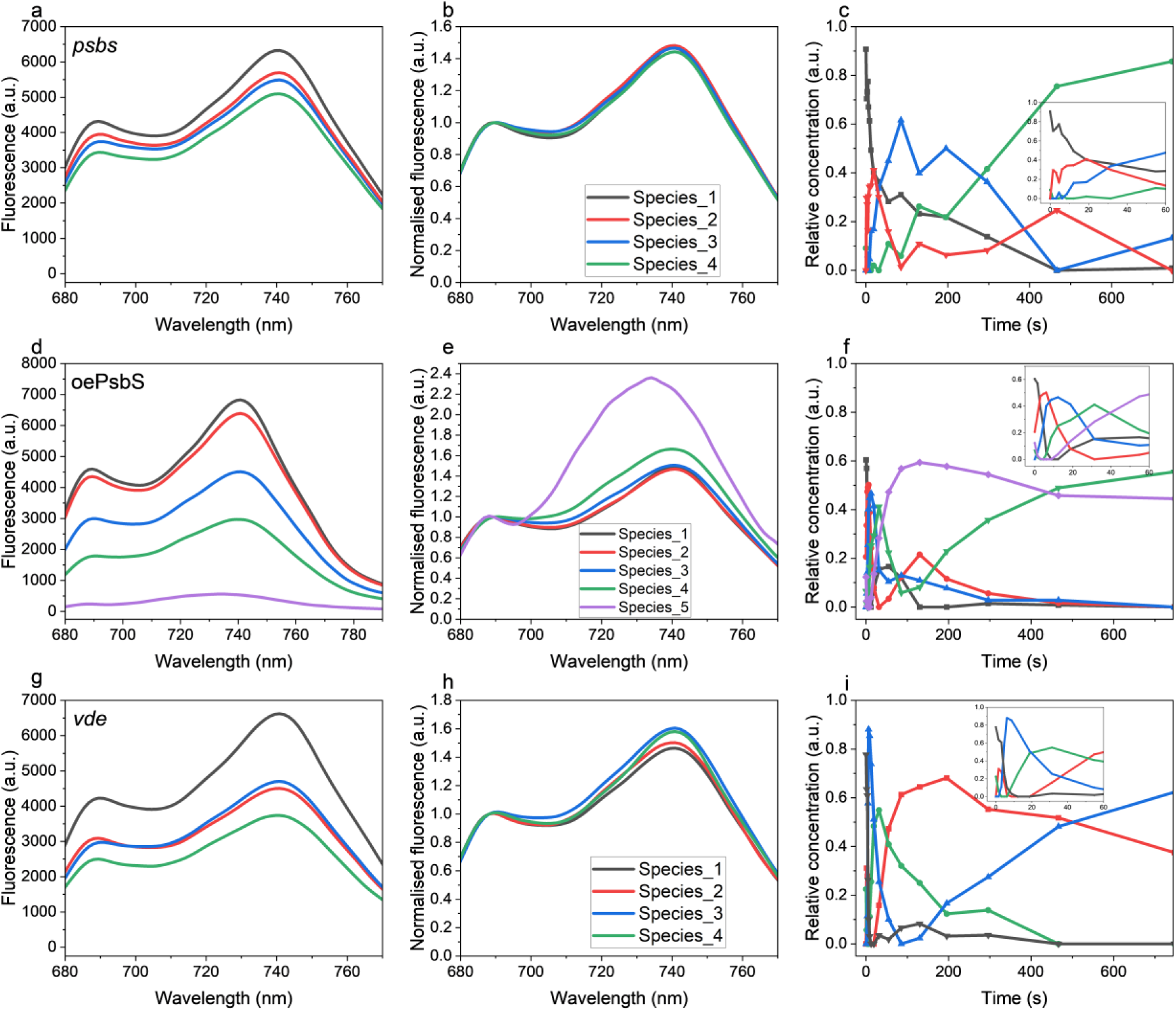
MCR-ALS analysis for hybrid aspen lines as SAES, normalised SAES at 688 nm and time dependent concentration profiles, respectively, for *psbS* (**a**, **b**, **c**); oePsbS (**d**, **e**, **f**) and *vde* (**g**, **h**, **i**). NPQ for all samples was induced by 600 µmol m^−2^ s^−1^ red actinic light.

Plants overexpressing PsbS (*Arabidopsis* L17, and particularly aspen oePsbS) showed similar kinetic development to WT plants, but all kinetic components were 2-4 times faster than in WT, which is consistent with the faster and stronger development of NPQ. Species 4 and, in particular, species 5 showed higher quenching in plants overexpressing PsbS (Figs. 2 and 3, Table 1). In *Arabidopsis* L17, the final ratio of the PSII-PSI component (species 5) to detached LHCII (species 4) was similar to WT, while oePsbS aspen had less PSII-PSI component but more detached LHCII than WT.

For *npq1* and *vde* aspen – affected in accumulation of Zea - four components/species were sufficient for a good fit. Species 2 and 3 (associated with LHCII detachment) accumulated but lacked a species similar to the PSII-PSI component (Figs. 2 and 3). The characteristic quenched LHCII (species 4) signature was also very faint. Therefore, we conclude that the lack of Zea prevents (or strongly delays) formation of the PSII-PSI component of NPQ. In contrast, *Arabidopsis npq2*, which also maintains high Zea levels in the dark, displayed similar kinetics to WT, displaying the fully developed PSII-PSI component (Fig. 2 j-l). In fact, formation of the PSII-PSI component was almost 10 times faster than in WT. Clearly, conditions for the formation of this component of quenching are already present in the dark, allowing very fast development of a quenched state.

### Light-induced changes in thylakoid stacking links to quenching

To study the thylakoid structure in intact leaves of the various *Arabidopsis* lines under dark-. (D) and actinic light-adapted conditions (hereafter the NPQ state) we used TEM. The actinic light treatment was the same as the one used to induce NPQ (Fig. 4a-d). The size of the grana stacks varied with illumination and to quantify the differences we measured both number of thylakoid layers per grana stack (Fig. 4e) and number of grana stacks per chloroplast (Fig. 4f). Fewer layers per grana stack were found in leaves in the NPQ state than in dark-adapted leaves: 24 % less in WT leaves (Fig. 4e), and 18, 18 and 11 % less in *npq1*, *npq4* & L17 leaves, respectively. Presence of Zea in the dark had a pronounced effect on the thylakoid structure. In D *npq2* leaves had almost as few layers per stack (4.4) as WT, *npq1* and *npq4* leaves in the NPQ state (3.9, 4.1 and 4.2, respectively) but this variable did not further decrease in the NPQ state. Reductions in numbers of layers per stack were associated with an increase in the number of stacks per chloroplast in WT & L17 plants (Fig. 4b). In WT leaves we observed 40 % more stacks in the NPQ state than in D, with a corresponding increase in L17 leaves of 11 %, but no significant changes in the numbers of stacks in *npq1*, *npq2* and *npq4* leaves.

**Fig. 4.**
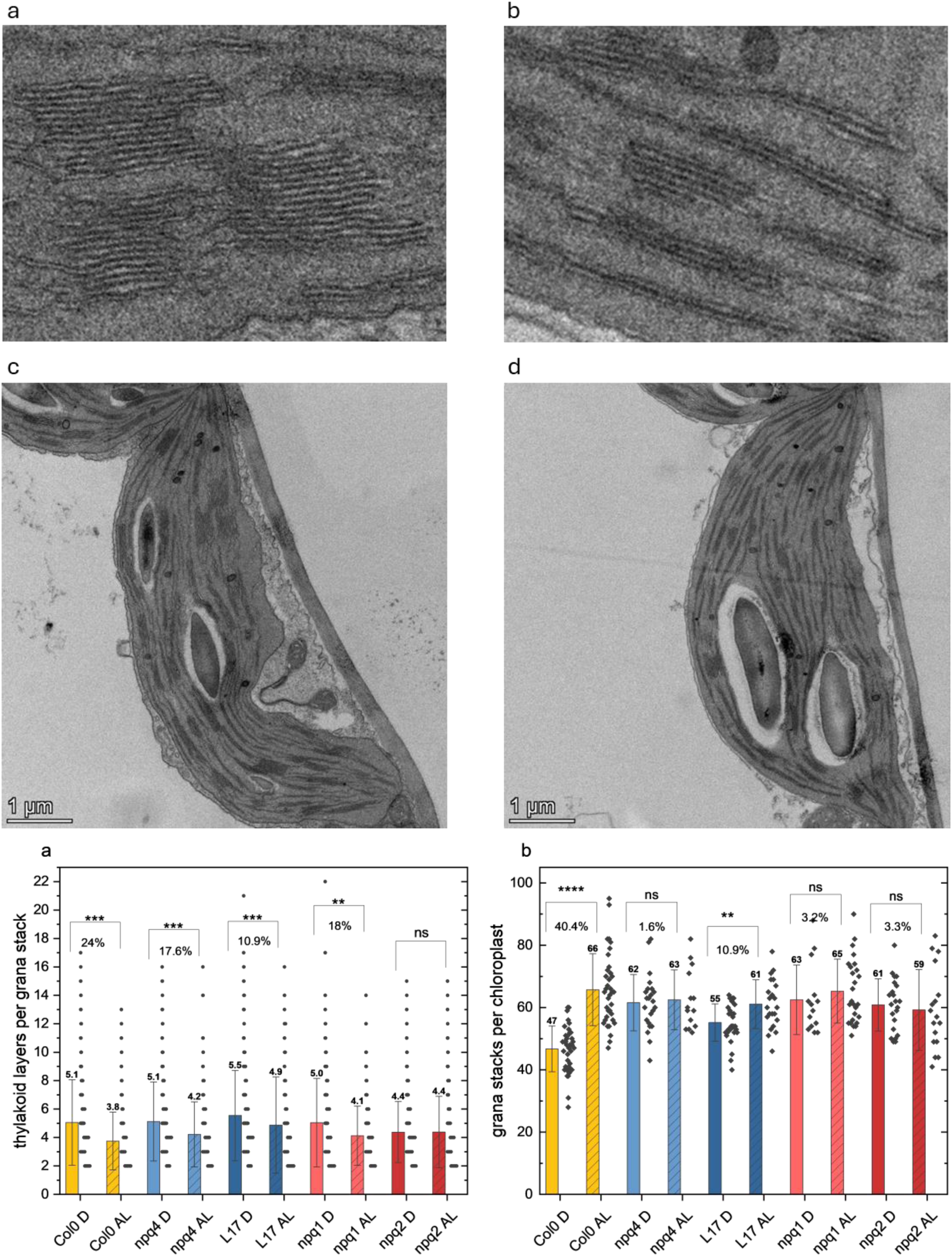
Magnified images of Col-0 in (**a**) D (night adapted, dark) and (**b**) AL (actinic light adapted/NPQ state, 600 µmol m^−2^ s^−1^) and the respective chloroplast images in (**c**) and (**d**). Thylakoid reorganisation upon NPQ induction measured as (**e**) number of thylakoid layers per grana stack and (**f**) number of grana stacks per chloroplast in *Arabidopsis* Col-0, *npq1*, *npq2*, *npq4* & L17 in D (solid bars) and AL (striped bars). Error bars represent standard deviation and mean values are represented above error bars calculated from n= 261 (Col0 D), 268 (Col0 AL), 276 (*npq1* D), 297 (*npq1* AL), 543 (*npq2* D), 296 (*npq2* AL), 255 (*npq4* D), 212 (*npq4* AL), 371 (L17 D), 200 (L17 AL) of grana stacks in (a) and n= 38 (Col0 D), 36 (Col0 AL), 14 (*npq1* D), 26 (*npq1* AL), 22 (*npq2* D), 17 (*npq2* AL), 23 (*npq4* D), 14 (*npq4* AL), 31 (L17 D), 22 (L17 AL) of chloroplasts in (b). Mann-Whitney test results (“*” = p < 0.05, “ns” = p ≥ 0.05) are represented for comparisons between D and AL for each genotype. Percentage change from D to AL conditions are represented below significance test results.

As the observed changes in thylakoid structure were complex, they are illustrated in additional ways in Fig. 5. A frequency distribution plot of the number of layers per grana stack shows how the proportion of layers per stack changed upon NPQ induction. Fig. 5a shows a comparison of the D and NPQ states of the various genotypes. Note, in particular, the strong shift towards two-layered stacks in WT and L17 leaves, from 20 to 35 and 11 to 29 grana stacks, respectively. The proportion of two-layered stacks also increased in *npq2* and *npq4* but not in the case of *npq1*. The increase in two-layered stacks in *npq2* came at the expense of three- and four-layered stacks.

**Fig. 5.**
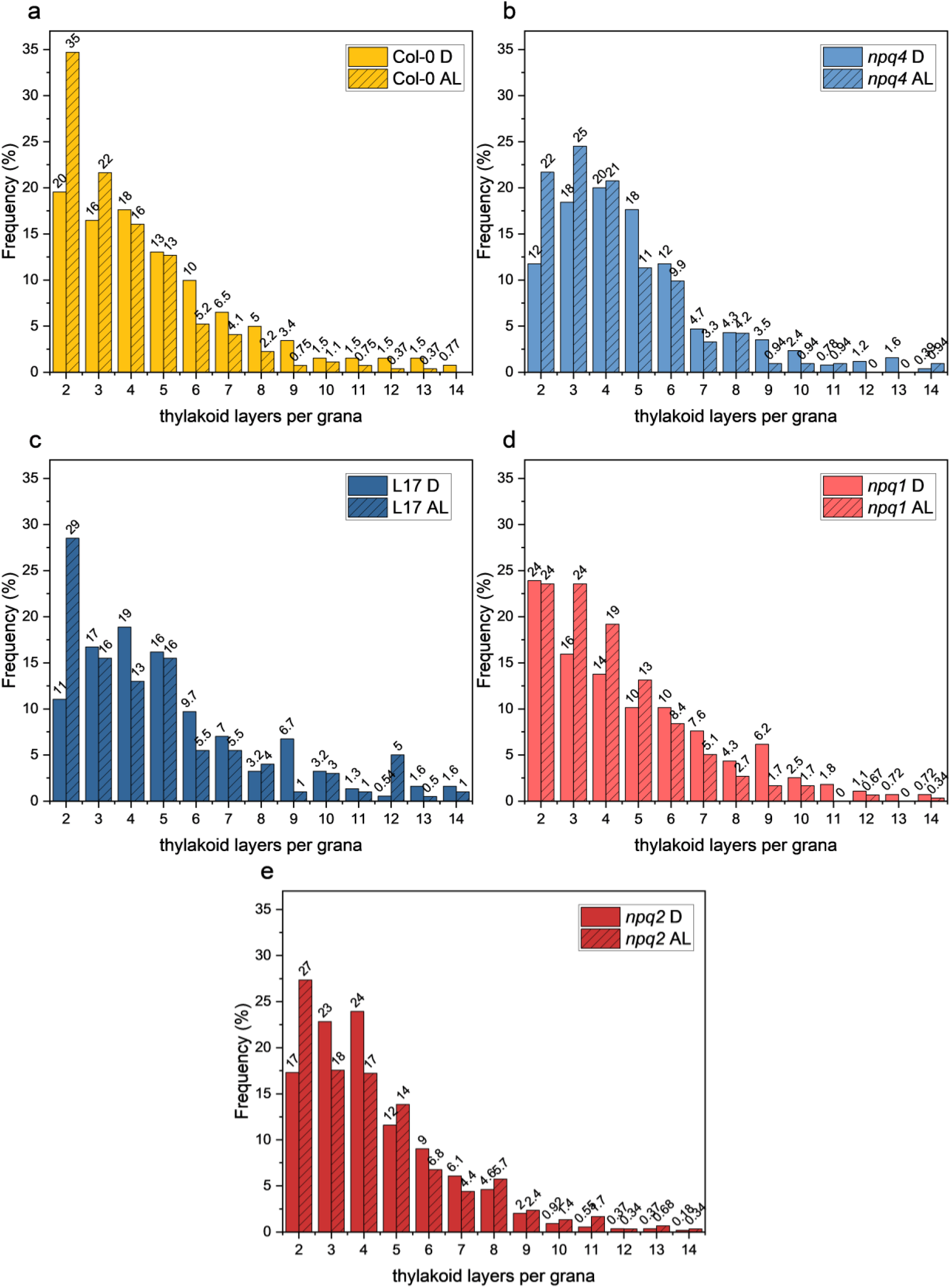
Frequency distribution of thylakoid layers per grana stack in *Arabidopsis* (a) Col-0, (b) *npq4*, (c) L17, (d) *npq1* & (e) *npq2* in D (solid bars) & AL (striped bars) conditions as measured in Fig. 4.

Our data show that both light conditions and the amount of Zea substantially influence thylakoid ultrastructure in a complex manner. Overall, induction of NPQ reduced grana stacking and 2-layered grana became more abundant. Thylakoids with excessive amounts of Zea (*npq2*) were the least stacked in D and light did not affect the degree of stacking which is consistent with the rapid development of NPQ in the *npq2* mutant, requiring only minor thylakoid rearrangements. While overexpression of PsbS slightly increased stacking, destacking in the NPQ state was less prominent.

## Discussion

Although NPQ mechanisms in angiosperms have been intensely studied in recent decades, detailed roles of the central molecular players - PsbS and Zea - and the relationships between qE and other putative mechanisms, such as qZ, qH and qI, are still not clear. We also do not understand the details about the relationships between mechanisms involving these players and the light-induced reorganization of thylakoid structure.

To probe these relationships, we combined simultaneous time- and spectrally-resolved detection of Chl fluorescence with global data analysis techniques to resolve the experimental NPQ kinetics into spectrally and kinetically distinct components, which we tentatively assign to biochemical entities. Application of this approach to *Arabidopsis* and aspen mutants with NPQ impairments, together with TEM structural analysis of thylakoids, provided illuminating indications of the key players’ roles, associated quenching mechanisms and links to thylakoid structural rearrangements. We acknowledge the fact that this analysis is complex, MCR-ALS has not previously been used to analyse this kind of data and the assignments of spectral components and the model might be questioned. But the analysis is robust in the sense that repeated measurements give similar results, and our interpretations are made considering the whole dataset (three plant species and mutants in key proteins). We also take into account previous data from for example ultra-fast spectroscopy of intact systems or isolated components. Our interpretations fit experimental data best and our proposed model is consistent. For example, the “branched model” for wild type plants where one species is split into two is needed for a best fit, not the alternative where the five components are populated one by one in sequence. Furthermore, components missing in plants lacking or overexpressing PsbS, or lacking VDE and ZE support our assignments. Future analysis with refined techniques will hopefully corroborate this model.

Self-absorption could complicate interpretation of fluorescence data. However, during the time course of our experiments we believe that there should be negligible changes in the optical properties of the leaf that could affect self-absorption. Therefore, as we in our analysis only compare the spectra with those of the unquenched states in the same leaf with the same self-absorption characteristics, they do not need to be corrected for self-absorption.

### PSII-PSI spectral component (species 5) indicates spillover

We interpret the transition from species 1 (unquenched PSII supercomplex, with weak unresolved PSI fluorescence) to species 2 as the partial or full detachment of, still unquenched, LHCII from PSII (Fig. 1j). In addition, energization of the membrane (which may cause very rapid spectral changes due to electrochromic shifts and fast light-induced changes (Sipka *et al*., 2022)) could contribute. Species 2 already provided half of the fluorescence at 10-20 s, indicating an almost total transient detachment of LHCII from PSII supercomplex. Species 3 corresponds predominantly to PSII cores. Over time species 2 decayed to the detached and quenched LHCII (species 4) and the highly quenched PSII-PSI component (species 5). Based on the dual nature of the spectra of species 5 we believe it to be a spillover complex, characterized by direct energy transfer from PSII to PSI.

The general picture – in terms of both relative populations and quenching ratios – is consistent with results of ultrafast fluorescence kinetic analysis of pine needles (Bag *et al*., 2020). In pine spillover seems to be a slightly larger component of NPQ compared to in *Arabidopsis* and aspen; the SAES of species 5 has a marginally larger far-red contribution. This is consistent with the higher maximal NPQ values in these samples (4.3 in pine *vs.* 2.3 in *Arabidopsis*). It could represent a species difference, but could also be a consequence of the prehistory of the material, plants grown under natural fluctuating light conditions (like our pine plants) show higher NPQ also at relatively low AL intensities (Schumann *et al*., 2017; von Bismarck *et al*., 2023).

We propose that spillover is the main light-induced quenching mechanism under ‘normal conditions’ also in *Arabidopsis* and aspen. Furthermore, fast and efficient development of the ‘spillover quenching’ state requires both PsbS and Zea, mutants lacking them do not develop a significant amount of the state and thus only small amounts of quenching. We observed strong quenching by spillover in *Arabidopsis*, aspen and pine. Therefore, we believe that spillover is a key quenching mechanism in all angiosperms and at least some gymnosperms. The resulting photoprotective effect of spillover quenching on PSII is far higher than the effects estimated from conventional PAM measurements, which typically yield NPQ values of around 2.5 in WT genotypes. The data presented here suggest that the corresponding ‘quenching factor’ could be ca. 4.7. However, the MCR-ALS analysis indicates that in the most efficient spillover milieu (aspen oePsbS) the reduction in fluorescence is about 13.3 in terms of quenching factor. Moreover, since about 50% of the fluorescence in the SAES of the spillover complex in fact derives from PSI, rather than PSII, the real quenching factor of PSII by NPQ could be twice as high. This is the factor by which NPQ downregulates PSII activity and any damaging side reactions resulting from the PSII charge separation (Lambrev *et al*., 2012). Therefore, NPQ has a huge dynamic range, underestimated by conventional PAM analyses but probably crucial for plants under highly fluctuating natural light conditions.

In an attempt to combine a significant amount of spillover with what is previously known about NPQ we have created a tentative model integrating quenching mechanism’s development, roles of PsbS and Zea in it, and associated structural arrangements that occur during NPQ induction. Functional and structural aspects of the model are illustrated in Figs. 6a and 6b, respectively. In the relaxed dark state most PSII is present in the form of super-complexes with attached LHCII, although smaller amounts of functionally detached, mostly unquenched, LHCII may be present. Switching on actinic light results in lumen acidification triggering NPQ. In WT leaves NPQ evolves over about 15 minutes but the fastest reactions – already leading to quenching - occur on the seconds timescale with pronounced changes in the fluorescence spectra. This fast reaction is a functional detachment of most LHCII complexes from the supercomplexes and subsequent pronounced quenching (with a similar time scale) of detached LHCII. The detachment and quenching processes are separate steps that can be resolved in *Arabidopsis*, but in aspen and pine they are faster and therefore more difficult to resolve. These fast quenching steps are strictly dependent on PsbS and represent what is termed “qE-quenching” in the literature.

**Fig. 6.**
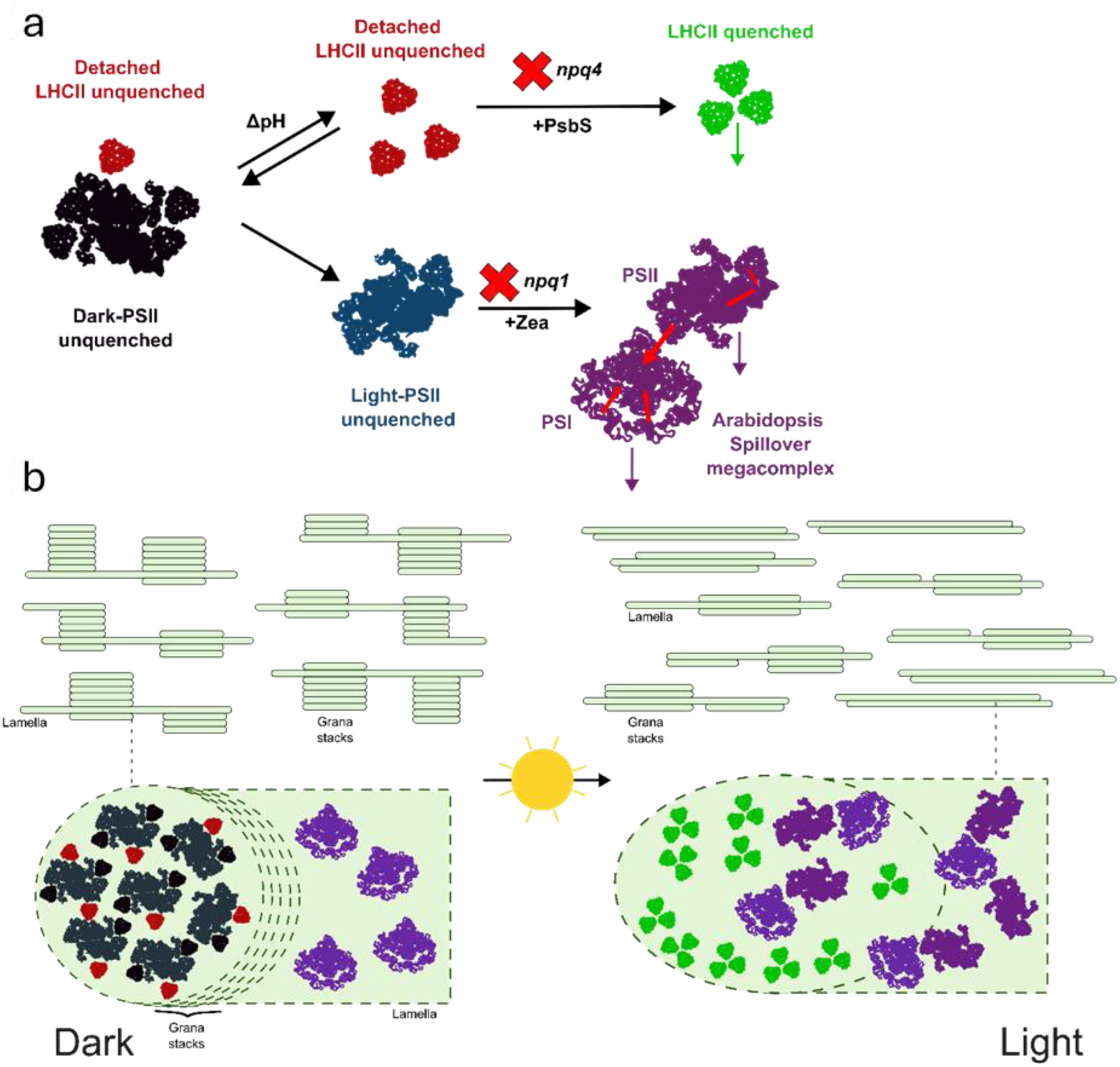
Model describing the proposed NPQ mechanism in plants. (**a**) During the dark-to-light transition, the pre-existing detached LHCII fraction (red) is enriched by the detachment of LHCII bound to the PSII core (black) by lumen acidification. In the presence of PsbS, the detached fraction rapidly switches to a quenched state (green), producing new Chl-Chl interactions which most likely quench excitation through charge transfer processes (green arrow). By the help of zeaxanthin, the reduced absorption cross-section PSII (blue) physically interacts with PSI producing the spillover megacomplex (purple). Red arrows represent hypothetical EET pathways between PSII and PSI, as well as between PSII and PSI antenna and their respective reaction centres. Purple arrows show decay pathways associated to PSII and PSI photochemistry. Red crosses indicate that the respective reaction step is drastically slowed down or not happening in the indicated mutant. (**b**) At a structural level, to increase the surface contact between PSI and PSII, the pronounced lateral heterogeneity present in the dark state is strongly reduced by a rearrangement of the thylakoid architecture. The left-hand side (dark) schematically shows the arrangement of a dark-adapted grana whereas right-hand section represents a fully light-adapted reorganized grana with reduced stacking. In the light-adapted state the thylakoid landscape is mostly dominated by the presence of spillover complexes and quenched LHCII complexes.

This actual photoprotection of PSII is not due to a real PSII non-photochemical excited state quenching but the very fast reduction of antenna size. LHCII-stripping of the PSII complex reduces the fluorescence yield, but this is merely due to a well-understood shortening of the excited state lifetime associated with a very fast reduction in antenna size rather than a quenching process *per se*. The actual PsbS-dependent quenching affects only the already detached LHCII and protects it from damage. This interpretation of these two fast processes resolved by MCR-ALS analysis is consistent with previous characterization of the PsbS-dependent quenching in the quasi-steady states by ultrafast time-resolved fluorescence analysis (Holzwarth *et al*., 2009; Bag *et al*., 2020). The underlying PsbS-dependent quenching mechanism is a Chl-Chl charge transfer (CT) mechanism and has been characterized both *in vitro* (Miloslavina *et al*., 2008; Ostroumov *et al*., 2020) and in proteoliposomes co-reconstituted with LHCII and PsbS (Pawlak *et al*., 2020). This mechanism gives rise to the characteristic far-red enhanced spectrum due to the charge recombination emission from the induced Chl-Chl CT state. In this study we have used these characteristic spectral features, along with the kinetic development (spectra and concentrations) that we can resolve, for the biochemical assignment of the underlying species. Obviously, assignments - like this - based mainly on spectroscopic features should be corroborated by biochemical studies that may give additional information, for example if the spectral components/species really have one-to-one relationships to biochemical entities.

After these fast steps most NPQ develops on a significantly longer time scale (about 50-100 s) in WT leaves. The species resulting from these slower steps is the most strongly quenched in the NPQ state. As argued above, its spectral features characterize this SAES component as a combination of PSII and pronounced PSI fluorescence developing concomitantly with the same kinetics. As this spectral feature – with both PSII and PSI contributions – develops with homogeneous kinetics it cannot merely be interpreted as a mixture of unconnected PSII and PSI particles. Instead, these features show that the underlying complex is a homogeneous coupled entity that emits both PSI and PSII fluorescence, the definition of a functional ‘spillover complex’. In this complex, energy is strongly quenched by energy transfer from PSII to PSI. The kinetics and spectral features of this complex in the NPQ state of pine have been characterized by ultrafast time-resolved fluorescence (Bag *et al*., 2020). Here we extend the findings to two angiosperm species.

Formation of the spillover complex is prevented by the absence of Zea and is largely hindered by the absence of PsbS. Therefore, PsbS has two roles in quenching regulation, acting as a direct inducer of LHCII quenching and participating in the spillover development (Fig. 6a). The connection of spillover complex formation with Zea and PsbS illuminates the associated structural requirements. Formation of a spillover complex involving coupling of PSII to PSI complexes requires a major thylakoid structural rearrangement. The pronounced lateral segregation of grana structures carrying PSII and stroma-regions carrying PSI is partly converted to a mixed structure with much less grana stacking (Fig. 6b). This is consistent with the structural arrangements during the dark-to-light transition for WT and mutants (Fig. S15).It is interesting that on long time scales (more than 300-500 s) some slow regulation occurs (cf. Figs. 1-3) in many genotypes as it leads to partial further attachment (sometimes also detachment) of quenched LHCII to the spillover complex, which may play a role in fine regulation of the quenching.

Presence of Zea in the dark dramatically accelerates spillover complex formation after exposure to light and strengthens quenching. Overall, the large majority of PSII complexes are quenched in these conditions, only a small fraction may remain in the unquenched state. However, the presence of Zea in *npq2* leaves does not lead to strong quenching in the dark-adapted state. This clearly conflicts with models suggesting that Zea plays a direct quenching role, for example in LHCII(Wilk *et al*., 2013; Sacharz *et al*., 2017), and even high amounts in the membrane may confer very little quenching capacity. In addition, LHCII isolated from *npq2* leaves (containing high levels of Zea) is only slightly quenched (with ca. 10-20 % shorter lifetime (Miloslavina *et al*., 2008)), far too little to explain huge *in vivo* NPQ effects. Furthermore, an NPQ value and strongly quenched spillover complex very similar to WT is induced in *npq2* leaves upon light activation (Fig. 2). This suggests that Zea in itself is not a strong quencher but could have a role in NPQ through involvement in the regulation of thylakoid structure, perhaps by reduce grana stacking.

### Grana destacking coincides with PSII-PSI spillover and is modulated by zeaxanthin during NPQ

Dynamic rearrangement of the thylakoid membranes is important for photosynthetic regulation and the extent of grana stacking varies with light conditions (Johnson and Wientjes, 2020). Recent evidence indicates that pronounced thylakoid lateral segregation is only present in dark-adapted leaves and under extreme conditions, like those of conifers in the boreal early spring (Bag *et al*., 2020)., most thylakoids may be in two layers, but even in milder conditions a significant fraction could be in thylakoid doublets where PSII and PSI centres could interact within spillover complexes (Fig. 6b). Robust evidence for the formation of PSII-PSI spillover-complexes, both *in vivo* and *in vitro*, has accumulated (Suorsa *et al*., 2015; Yokono *et al*., 2019; Bag *et al*., 2020; Yokono, Ueno and Akimoto, 2021; Kim *et al*., 2023; Terashima *et al*., 2024; Yokono, Noda and Minagawa, 2024) and we have exposed mutants in key proteins for NPQ to various light conditions and noted changes that by large corroborate earlier findingsIt has been speculated that stacked doublets or ‘hybrid G-SL regions’ form in response to selective pressures to promote linear over cyclic electron transport (Garty *et al*., 2024) but we propose that the capacity to increase NPQ through spillover is a more important driver. The mechanism and structural requirements of cyclic electron flow around PS I are not well understood, but NPQ (Niu *et al*., 2023) and spillover complexes may also play a role in that mechanism.

The observed changes in thylakoid structure upon NPQ induction highlight the complex relationship between grana stacking and Zea. Our data show that NPQ induction generally leads to a reduction in grana stacking, with an increase in the proportion of two-layered stacks. In the *npq2* mutant, which accumulates excess Zea, the mean number of layers per grana stack was lower in the dark-adapted state (4.4 in *npq2* compared to 5.1 in WT), and light exposure did not further reduce stacking (Fig. 4). Notably, the *npq1* mutant showed no shift towards two-layered stacks, in contrast to WT and L17 plants, where two-layered stacks increased significantly upon NPQ induction, and to *npq2* and *npq4* mutants, where the increase was less pronounced (Fig. 5). However, *npq2* plants showed an increase in two-layered stacks at the expense of three- and four-layered stacks (Fig. 5e), supporting the idea that thylakoid rearrangement by destacking is essential for facilitating PSII-PSI spillover. Grana destacking and the increase in two-layered thylakoids may create greater opportunities for PSII-PSI spillover within the grana end membranes and at grana margins. The presence of two-layered stacks increases the surface area for potential spillover at the grana end membranes. Alternatively, spillover may occur between a PSI complex, located in the grana end membrane or at the margins, and a PSII complex in the adjacent appressed membrane. These observations suggest that Zea plays a crucial role in the structural rearrangements required for the rapid development of NPQ, as proposed in the NPQ model (Fig. 6).

## Conclusion

To conclude, we suggest a tentative model for NPQ integrating the functions of PsbS, Zea, spillover complexes and thylakoid rearrangements. However, it does not explain what forces hold the spillover complex together or how thylakoid remodelling is triggered. The interactions between PsbS, Zea, light activation and thylakoid stacking are complex, and we have fragmentary understanding of the regulatory and structural forces controlling and shaping thylakoid rearrangement and spillover complex formation. Thylakoid protein phosphorylation is probably involved and previous suggestions that rearrangements are related to the balance between linear and cyclic electron flow (Garty *et al*., 2024) are not mutually exclusive to the formation of spillover complexes. We believe that spillover is the major regulatory component of NPQ under most ‘natural’ conditions, not only high and/or fluctuating light where cyclic electron flow appears to be most relevant. Under these conditions the lateral segregation of PSII and PSI in the thylakoid membrane may not be pronounced. We also suggest that many currently used techniques underestimate the extent that NPQ-processes can provide photoprotection. Our findings could be relevant for efforts to increase crop yields by modifying NPQ (Kromdijk *et al*., 2016; De Souza *et al*., 2022).

## Methods

### Construction of hybrid aspen mutants

We made CRISPR constructs to create T89 hybrid aspen (*Populus tremula* x *Populus tremuloides*) mutants with knocked-out *PsbS* and *VDE* genes, hereafter *psbs* and *vde* mutants. The *PsbS* and *VDE* genes in aspen were identified based on amino acid sequence similarity to their homologs in *Arabidopsis thaliana*. A pair of sgRNAs for each target gene were identified with CRISPR-P 2.0 (http://crispr.hzau.edu.cn/CRISPR2/) (Table S1). Potential off-target sites for sgRNAs were also checked against the hybrid aspen T89 genome using the BLAST methodology implemented in PlantGenIE (https://plantgenie.org/BLAST). sgRNAs were introduced into entry vectors by site-directed mutagenesis PCR. The final vector (containing promoter, Cas9 CDS, terminator, the two sgRNAs and an antibiotic resistance cassette) was assembled by GreenGate reactions as previously described (Lampropoulos *et al*., 2013; André *et al*., 2022). PsbS-overexpressing (hereafter oePsbS) hybrid aspen lines were constructed by introducing the PsbS coding sequence from *Populus tremula* (Fig. S1) under control of the 35S promoter in the pk2gw7 destination vector using GATEWAY protocols.

*Escherichia coli* strain DH5α was used for amplification of all plasmids, and their identities were confirmed by sequencing. Final vectors were transferred into hybrid aspen T89 using a standard procedure (Nilsson *et al*., 1992). Transgenic lines from each transformation were screened for target gene deletions by PCR with primers listed in Table S1. The images and legends of Figs. S2 and S3 present and discuss representative genotyping results for the lines used in the study reported here and subsequent sequencing results. PsbS protein expression in overexpressing and mutant lines was tested by immunoblotting (Fig. S4). Several *psbs*, *vde* and oePsbS lines were generated and analysed. One line with each modification was chosen for the detailed analysis presented in this article.

### Plant material & growth conditions

Leaves of 6-week-old *Arabidopsis thaliana* Col-0 (wild type), *npq1*, *npq2*, *npq4* and L17 plants grown in the greenhouse (with 150-180 µmol m^−2^ s^−1^ growth light) in short day conditions (8 hr light, 22 °C/16 hr dark, 18 °C cycles) were used for measurements. Leaves of 6-to 7-week-old hybrid aspen T89 and mutant lines were harvested from trees grown in long day conditions (16 hr light, 22^0^C/8 hr dark, 18^0^C cycles) in the greenhouse (with 70-90 µmol m^−2^ s^−1^ growth light). Pine (*Pinus sylvestris*) needles were sampled from a tree, at least 40 years old, grown outdoors in August 2023 (when the mean temperature was 16 °C). The needles were arranged to form a continuous surface for chlorophyll fluorescence measurements.

### Chlorophyll fluorescence analysis

Chlorophyll fluorescence measurements were obtained using a ChloroSpec L1 *in vivo* fluorescence spectrometer (ChloroSpec B.V. Amsterdam, www.chlorospec.com (Nanda *et al*., 2024). All samples were dark-adapted for 30 min before measurements at room temperature unless mentioned otherwise. A burst of single turnover flashes (STF) (250 pulses, each 130 µs wide, separated by 3 ms intervals) of 60,000 µmol m^−2^ s^−1^ red excitation light was used as saturating light for the NPQ sampling at indicated times, starting with a dark-adapted sample. This STF burst – sufficient to close all the PS II reaction centres – was immediately followed by a short (700-1000 µs MTF (multi-turnover flash) pulse of 15,000 µmol m^−2^ s^−1^ red excitation light, during which the time-resolved spectra used for the NPQ kinetic analysis were recorded. In all cases NPQ was induced using 600 µmol m^−2^ s^−1^ red actinic light (AL). Depending on the widely varying kinetics of the tested genotypes the NPQ actinic induction phase was measured over a time range of approx. 800-1000 sec, followed by a dark relaxation phase of approx. 700-800 sec. Corresponding data are shown in Fig. S6 for the *Arabidopsis* genotypes and pine needles as plots of the NPQ values at three selected wavelengths. Fig. S7 shows the full NPQ spectra at indicated actinic/relaxation times. As previously reported (Nanda *et al*., 2024), these figures demonstrate the pronounced emission wavelength dependence of the NPQ values, as well as the pronounced genotype dependence. Generally, NPQ is maximal at around 680 nm and minimal at around 720 nm. Note the very large NPQ values, around 5, for both pine and *Arabidopsis* L17 plants. Corresponding data for the hybrid aspen genotypes are shown in Figs. S8 (NPQ kinetics) and S9 (NPQ spectra). They show similar pronounced wavelength dependence of NPQ to the *Arabidopsis* and pine samples. Here we focus on the kinetics of the NPQ induction, rather than the relaxation, phase. This is because previous work (Niu *et al*., 2023) and our data suggest that hysteresis effects do occur in the relaxation phase, which thus requires more complex analysis.

### Spectro-kinetic data analysis

The time-resolved fluorescence spectra (3D intensity-wavelength-time surfaces) recorded during the NPQ induction phase were decomposed into their species-associated emission spectra (SAES) and time-dependent concentration profiles of the various components in an iterative process using multi-component spectro-kinetic analysis. For reasons previously discussed (Holzwarth, Lenk and Jahns, 2013) the kinetic analyses of the NPQ involved analyses of changes in fluorescence intensity with time, rather than changes in NPQ *per se* with time. One analytical method to decompose time-resolved NPQ spectra – global target modelling (Holzwarth, 1996) – has been previously introduced (Nanda *et al*., 2024). This modelling technique decomposes time-and wavelength-resolved three-dimensional spectral surfaces using a kinetic model described by a set of (pseudo-)first-order kinetic differential equations. While this method works well over short time-ranges of NPQ induction (up to ca. 200 s, depending on the conditions), kinetic target modelling proved to be unsuitable for describing the NPQ kinetics over the entire time range of interest in our NPQ induction experiments (ca. 1000 s of induction). This is because the experimental fluorescence kinetic traces display features (bumps and other complex long-time trends and behaviours) that cannot be described by first-order kinetic models. Moreover, kinetic target modelling also has limitations for correctly describing reversible reactions. Therefore, we opted for a less restrictive method to deconvolute the spectro-kinetic data into their components: multivariate curve resolution (MCR) in combination with an alternating least squares (ALS) algorithm. MCR-ALS has well established utility for spectro-kinetic analysis of complex chemical systems (Tauler, 2007; Ruckebusch and Blanchet, 2013). MCR-ALS results are the same as those of kinetic target modelling: dissection of the experimental data into components by separating them into their spectral (SAES) and time-space (concentration developments in time) components. A major advantage of MCR-ALS is that it does not require assumptions about an unknown kinetic model and starting concentrations of the components. On the other hand, MCR-ALS has mathematical restrictions that must be fulfilled to exclude ambiguities and solutions that are not physically reasonable. These are: fulfilment of a bilinear form, non-negativity constraints for concentration profiles and spectral forms (SAES), and the ‘closure’ condition. In physical terms the mathematical term ‘closure’ equates to the mass balance conditions, here a constant sum of the concentrations profiles (Tauler, 2007; Abdollahi and Tauler, 2011). These conditions are fulfilled in NPQ experiments if the total Chl concentration is constant and the SAES of each component does not change during the experiments. Note that the ‘concentration profile’ does not mean the actual chemical concentrations of the species, but the total absorption cross-sections associated with the components and their respective SAES. Besides these fundamental restrictions and conditions, the ability to resolve a certain number of components is (as in all fittings) determined by the signal/noise ratio. To improve precision and efficiency in parameter estimation, while avoiding overfitting and ambiguities in the least-squares optimization algorithm a diagonal Tikhonov regularization term (Tikhonov *et al*., 1995; Voss, 2010) was added in the fitting procedure to the norm of the residual. The quality of fits was judged by residual plots, and fitting was considered good if the large majority of deviations did not exceed ±2σ.

Although not strictly required, and in most cases not crucial, our implementation of MCR-ALS typically uses starting guesses for the SAES, obtained from kinetic target analysis performed over a short time range (approx. 120 s). Results of these target analyses for the *Arabidopsis* and hybrid aspen lines are shown in Figs. S10-S11. The graphs to the left in these figures show the resulting SAES, and graphs to the right show the corresponding concentration profile and central graphs the normalized (at 688 nm) SAES that better display the spectral differences between SAES for subsequent assignment to actual biochemical species/complexes. Corresponding residual plots (showing the quality of fit) are provided in Fig. S12 and the residual plots for fitting over the full time ranges for MCR-ALS analyses in Fig. S13. In most cases four or five component spectra were required to properly describe the kinetics within the fitting criteria (deviation by > 2σ at only a few time points, at most, in the residual plots). However, in a few cases the first 2-3 s of the NPQ kinetics were not well fitted, by either target analysis or MCR-ALS. This was probably due to fast spectral changes resulting from energization of the membrane (Junge, 1977) and/or fast light-induced relaxations in the antenna/reaction centre complexes (Sipka *et al*., 2021, 2022). Adding more spectral components could improve fitting, but the signal-to-noise ratio of these measurements did not permit trustful decomposition into more than five (or sometimes six) components. In contrast to target analysis, MCR-ALS allows exclusion of these first 2-3 poorly fitting data points from the fitting, without affecting the fitting at later time points. This option was chosen when the early disturbance had a significant effect on the resulting SAES component spectra. Since the NPQ kinetics we describe all have ≥10 s halftimes, this did not affect our results.

The meaning and interpretation of the properties of the SAES and concentration profiles are the same in MCR-ALS analysis and target analysis (Nanda *et al*., 2024). The shape of each SAES is characteristic of the biochemical nature of the species while the area under the SAES is proportional to its the fluorescence yield. Since we did not measure absolute intensities in this study the areas under the curves (*i.e.*, amounts of quenching) are normalized to the area of the most populated (and unquenched) species in the dark-adapted state (Table 1). We applied multiple criteria for the assignment of a spectral form (SAES) to a biochemical species. One was its spectral shape, relative to the SAES of the unquenched starting (dark) species rather than the absolute spectral shape (or spectral change). This is because the absolute - but not the relative - shape of the SAES is influenced by leaf self-absorption. These ‘concentration profiles’ do not represent the actual concentrations, but rather concentrations weighted by the associated antenna sizes (essentially the relative amounts of Chls associated with a particular species). Nevertheless, it is appropriate to think in terms of ‘concentrations’ and we use this terminology throughout this paper. Finally, an SAES can be assigned a biochemical species – *inter alia* - by either comparison with the spectral shapes obtained in ultrafast time-resolved experiments (Holzwarth and Jahns, 2014; Bag *et al*., 2020; Pawlak *et al*., 2020) with leaves and isolated components, or the inspection and interpretation of the concentration profiles in a kinetic model.

### TEM analysis

Slices from the middle region of *Arabidopsis* leaves in night-adapted (16 hours dark) and AL-adapted (630 nm red light at 600 µmol m^−2^ s^−1^ intensity for 30 minutes) conditions were cut in distilled water and fixed in 4% paraformaldehyde and 2.5% glutaraldehyde in 0.1 M sodium cacodylate buffer, pH 7.4 (all from TAAB Laboratories, Aldermaston, England) for 1 hour in dark at room temperature. Samples were rinsed with sodium cacodylate buffer and incubated in 1% osmium tetroxide (TAAB Laboratories, Aldermaston, England) in dark for 2 hours at room temperature. The fixed material was dehydrated in increasing concentrations of ethanol and eventually embedded in Spurr resin (TAAB Laboratories, Aldermaston, England). A diamond knife was used to obtain 70 nm ultrathin sections that were contrasted in uranyl acetate and lead citrate. Images of the sections were acquired with a Talos 120 C electron microscope (FEI, Eindhoven, The Netherlands) operating at 120 kV in combination with a Ceta 16 M CCD camera (FEI, Eindhoven, The Netherlands). ImageJ software was used to obtain thylakoid parameters from chloroplast images. We counted mean number of thylakoid layers per grana stack in 200-543 grana stacks and number of grana stacks per chloroplast in 14-38 chloroplasts of each genotype subjected to each treatment. Representative chloroplast images of *Arabidopsis* lines used for analyses are shown in Fig. S14.

## Abbreviations

NPQ: Non-photochemical quenching
PSII: Photosystem II
PSI: Photosystem I
LHCII: Light Harvesting Complex II
PsbS: Photosystem II Subunit S
Zea: Zeaxanthin
VDE: Violaxanthin Epoxidase
ZE: Zeaxanthin Epoxidase
qE: Energy-dependent quenching
qZ: Zeaxanthin-dependent quenching
qH: Lipocalin-dependent quenching
qI: Photoinhibition-dependent quenching
Chl: Chlorophyll
PAM: Pulse-Amplitude Modulated
STF: Single-Turnover Flash
MTF: Multi-Turnover Flash
AL: Actinic Light
GTA: Global Target Analysis
MCR-ALS: Multivariate Curve Resolution - Alternating Least Squares
TEM: Transmission Electron Microscopy
SAES: Species-Associated Emission Spectra
WT: Wild Type
oePsbS: Overexpressing PsbS

## Acknowledgements

The presented research was supported by funding from the Swedish Research Council VR, Swedish Foundation for Strategic Research (SSF), Kempestiftelserna, Knut and Alice Wallenberg Foundation, Vinnova and Trees for the Future (T4F)

## Author contributions

SN, MC, TS, ARH and SJ planned the experiments. SN contributed to all experiments and analyses. MC and TS participated in the fluorescence and TEM analyses, VF and NF participated in the construction of aspen mutants and transgenics, JL performed pigment analyses and PB contributed to the interpretation of the results. SN, MC, ARH and SJ wrote the paper with contributions from all authors.

## Competing interests

ARH is a cofounder of ChloroSpec B.V.

## Notes

### Competing Interest Statement

Alfred R. Holzwarth is a cofounder of ChloroSpec B.V.

