## Supplemental File for "Spillover is the dominant non-photochemical quenching mechanism in angiosperms"

5’ → 3’
ATGGCTCAAACCATGGTGCTCATGTCTGGTGTCTCTACGAGGCAAGTGGTGGACTTGAAGAGAGACC
CTTTACTTCAGTTTCAAGTTCAGAGACTAAGGCCTGCACCATTCTCTCGTCTTCTCTACAATCCTCTTCC
TAGCAAAGCTTCTTCATCCAATGCTTTTACTACGCTTGCCCTCTTCAAGCCCAGAACCAAGGCGGTTC
CTAAGAAGGCTGCTCCACCACCGAAGCCAAAGGTTGAAGATGGTATTTTTGGTACCTCCGGTGGCAT
CGGTTTCACTAAGCAGAATGAGCTCTTCGTGGGACGTGTTGCCATGATTGGATTTGCTGCATCGTTGTT
GGGAGAGGCAATAACAGGAAAAGGAATTCTATCTCAGCTGAACCTAGAGACTGGAATTCCCATTTACG
AAGCAGAACCACTTCTTCTTTTCTTCATCCTTTTCACCTTGCTCGGAGCCATTGGAGCTTTGGGTGATC
GCGGCCGCTTCGTTGATGACCCACCAACCGGCATTGAAGGAGCTGTGATCCCTCCAGGAAAAAGCT
TGAGGGCAGCATTGGGTCTCAAGGAGGGAGGTCCTCTATTTGGATTTACAAAATCCAACGAACTTTTC
GTGGGGAGATTGGCTCAGCTGGGCATCGCTTTCTCTCTAATTGGAGAAATCATAACTGGAAAGGGAG
CTCTAGCACAACTCAACATCGAGACAGGGATTCCAGTCAGTGAAATCGAACCACTTGTGTTGTTCAAC
GTTCTCTTCTTCTTCGTTGCCGCATTGAATCCCGGGACCGGGAAATTCGTGACCGATGAGGATGAAGA
GTAG

**Fig. S1** Coding DNA sequence (CDS) sequence for PsbS in *Populus tremula* with gene ID Potra2n2c5662 (transcript 2).

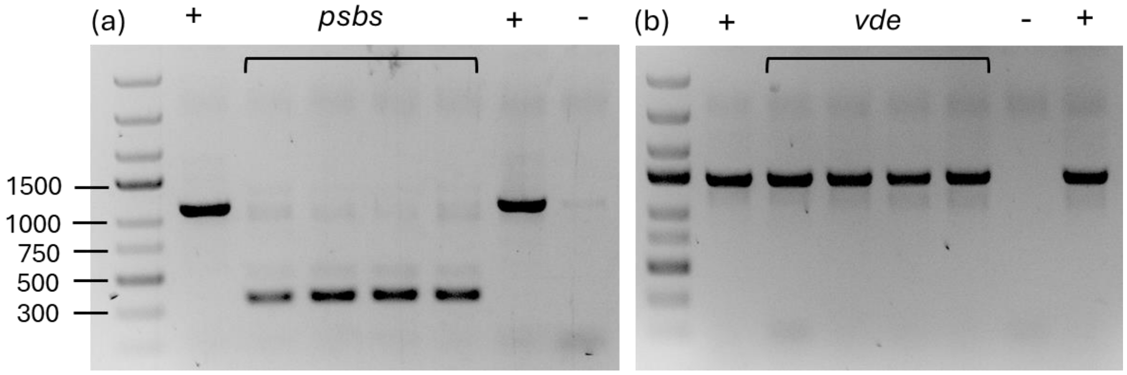


**Fig. S2** Gel electrophoresis after PCR with genotyping primers (c.f. Table S1) for 4 biological replicates of *psbs* & *vde* lines. (**a**) The *PsbS* gene is detected at 1200 bp in the wild-type background and with a deletion at 350 bp in the *psbs* lines indicating that it is a deletion mutant line. (**b**) *VDE* gene is detected at 1460 bp in the wild type and *vde* lines indicating that *vde* has a point mutation making the protein non-functional in the plant (c.f. S14) but is seen here as a wild-type band. The phenotype is verified as the NPQ induction is hindered in this line (Fig S4 (d)). “+” indicates wild-type T89 template and “-” indicates a control with no added template for the PCR reaction.

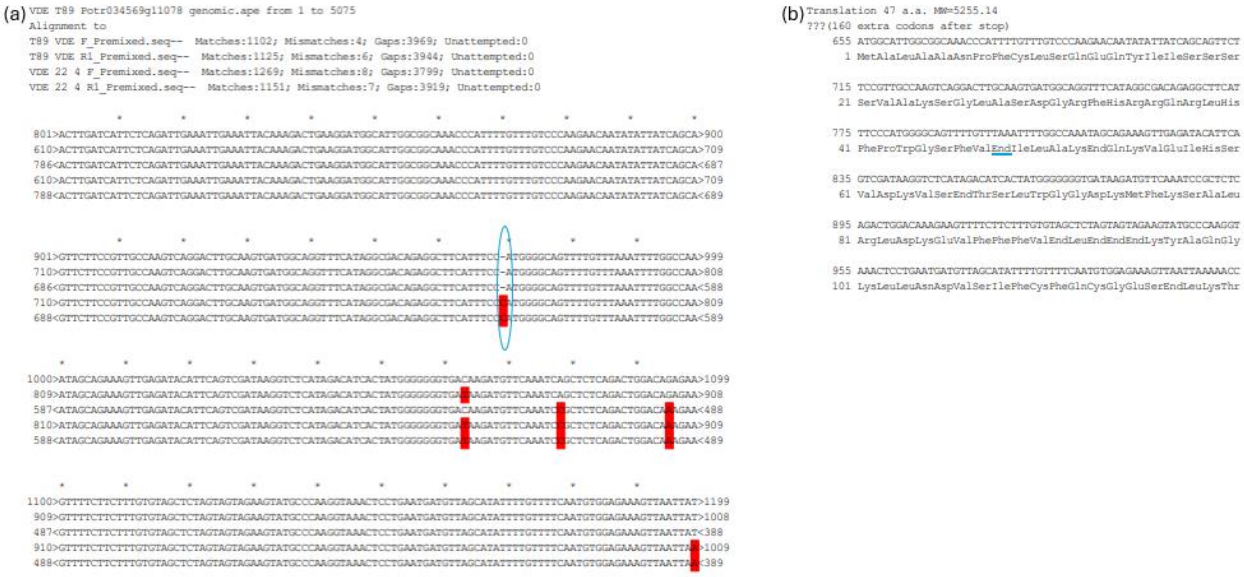

**Fig. S3** (**a**) Alignment of sequencing results of T89 wild type and *vde* PCR products (c.f. Fig S13) with *VDE* gene specific primers (c.f. Table S1) reveals a single base pair insertion (circled) in the mutant line which leads to an early stop codon (underlined) in the *vde* mutant line depicted in (**b**).

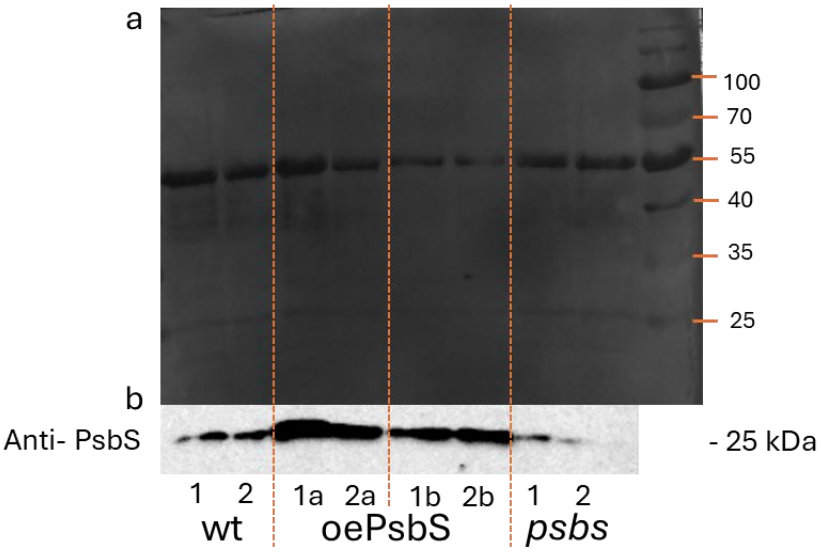


c

| Sample | CR250 stain at 55 kDa | Anti-PsbS band at 25kDa | PsbS /CR250 band ratio |
|---|---|---|---|
| wt 1 | 31377.45 | 15194.5 | 0.5 |
| wt 2 | 22004.31 | 25177.71 | 1.1 |
| oePsbS 1a | 26275.92 | 50627.77 | 1.9 |
| oePsbS 2a | 14123.99 | 38889.06 | 2.8 |
| oePsbS 1b | 11245.62 | 31569.99 | 2.8 |
| oePsbS 2b | 6962.53 | 37042.59 | 5.3 |
| *psbs* 1 | 19557.46 | 10709.02 | 0.5 |
| *psbs* 2 | 24854.66 | 2761.397 | 0.1 |

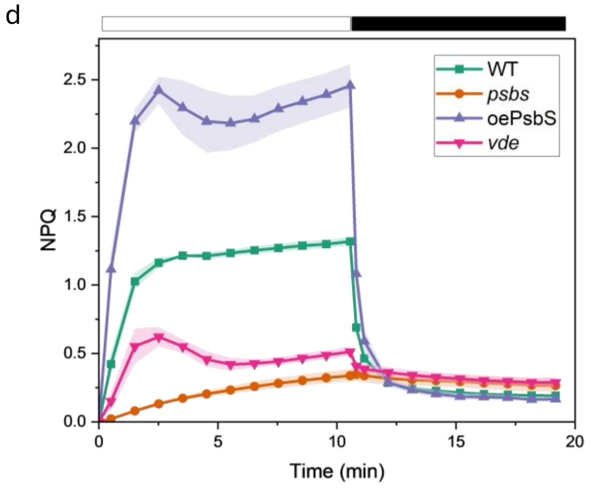


**Fig. S4** (**a**) Coomassie (CR250) stained gel image and (**b**) immunoblot membrane with anti-PsbS. 2 biological replicates are indicated by 1 & 2 for each line. 10 µl of total protein extract from 10mg leaf tissue was loaded for all samples. Relative quantification was done by normalising band intensity values of the immunoblot by Coomassie bands at 55 kDa in (**c**). NPQ kinetics plot obtained at ~730 nm by SpeedZen (imaging fluorescence instrument Johnson et al, 2009) for the corresponding lines and *vde* with 3 biological replicates is shown in (**d**). The NPQ was induced using 1000 µmol $m^{-2}$ $s^{-1}$ red actinic light.

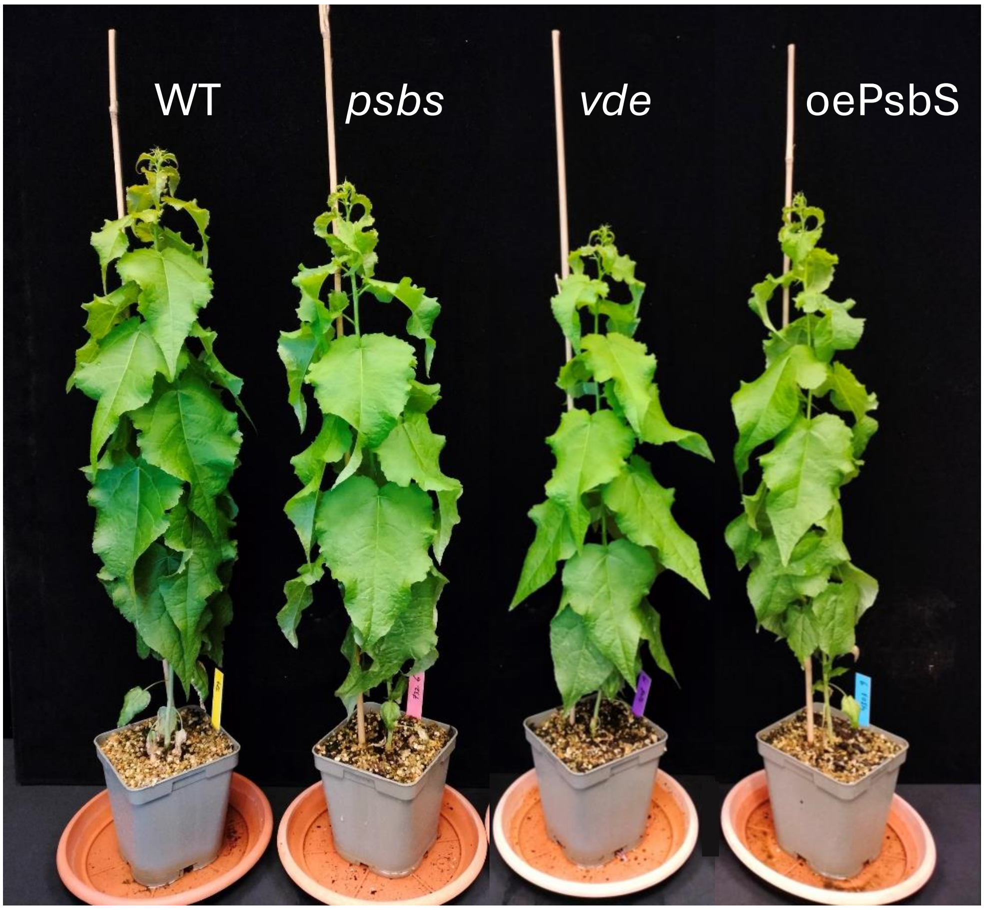


**Fig. S5** Representative images of 7-week-old aspen lines grown in the greenhouse where the growth light intensity ranged between 70 to 90 µmol $m^{-2}$ $s^{-1}$. The trees were grown in long day conditions where the day lasted for 16 hours at 22 ºC and the night lasted for 8 hours at 18 ºC.

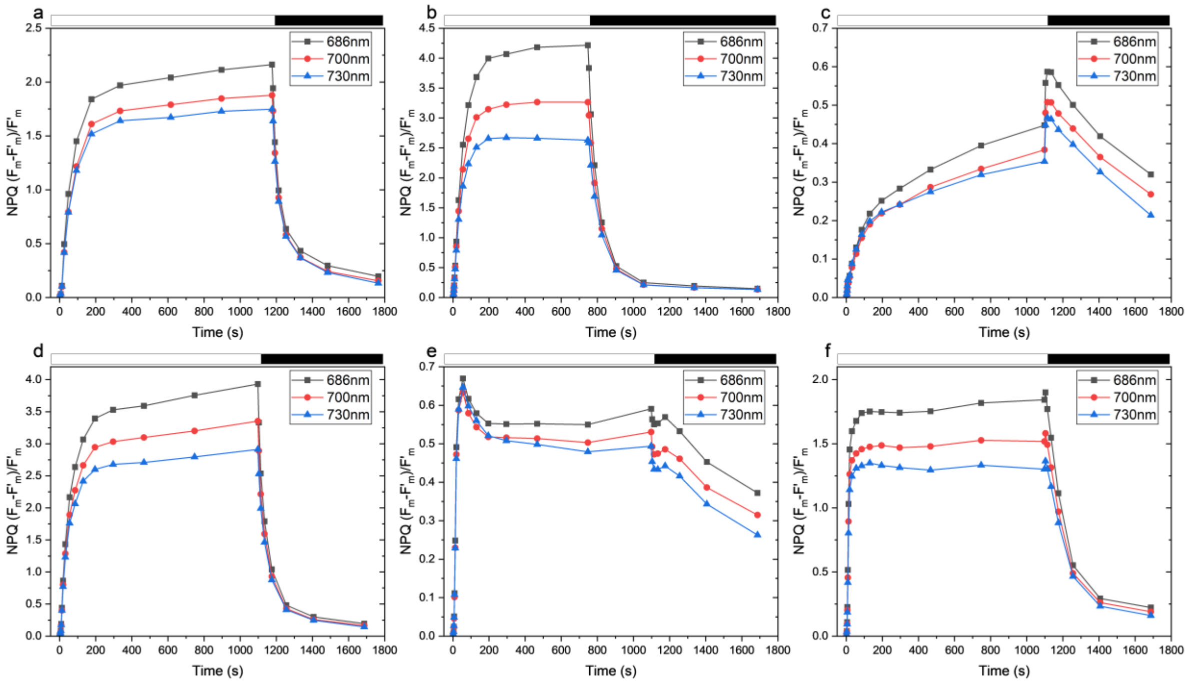


**Fig. S6** NPQ induction and relaxation kinetics of (**a**) *A. thaliana* wild type, (**b**) *P. sylvestris*, (**c**) *A. thaliana npq4*, (**d**) L17, (**e**) *npq1* and (**f**) *npq2*. All samples were induced with 600 µmol m$^{-2}$ s$^{-1}$ red actinic light.

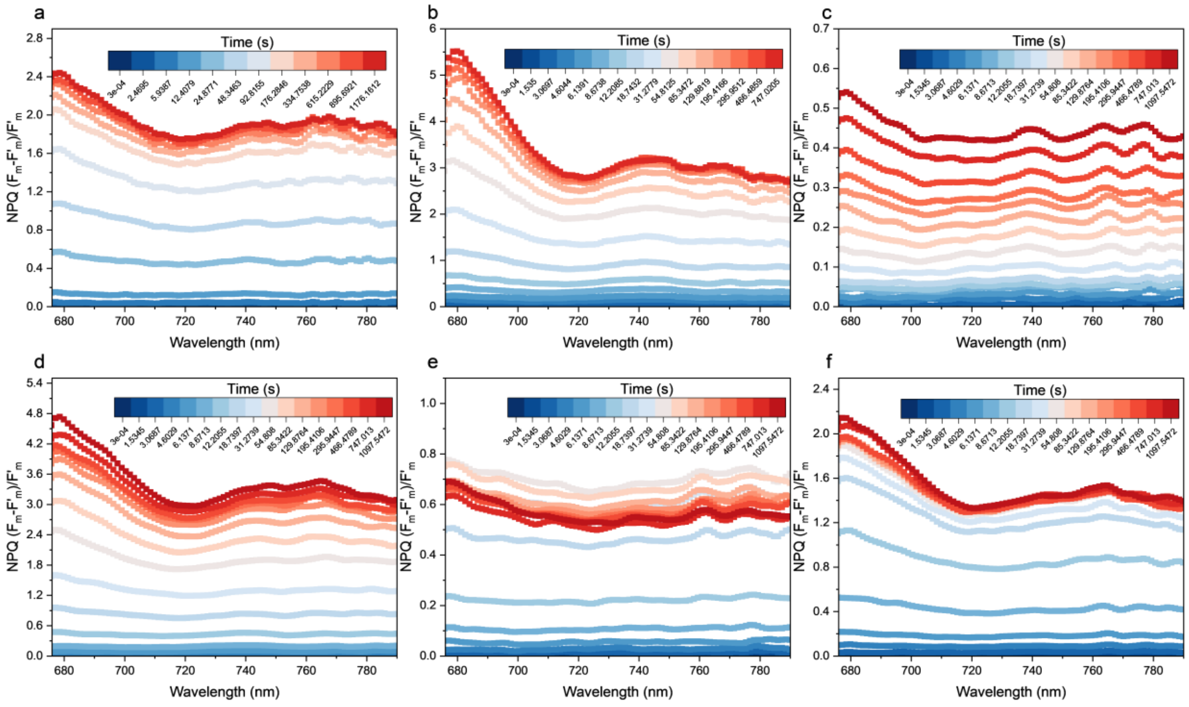


**Fig. S7** NPQ spectra of (**a**) *A. thaliana* wild type, (**b**) *P. sylvestris*, (**c**) *A. thaliana npq4*, (**d**) L17, (**e**) *npq1*and (**f**) *npq2*. All samples were induced with 600 µmol m$^{-2}$ s$^{-1}$ red actinic light.

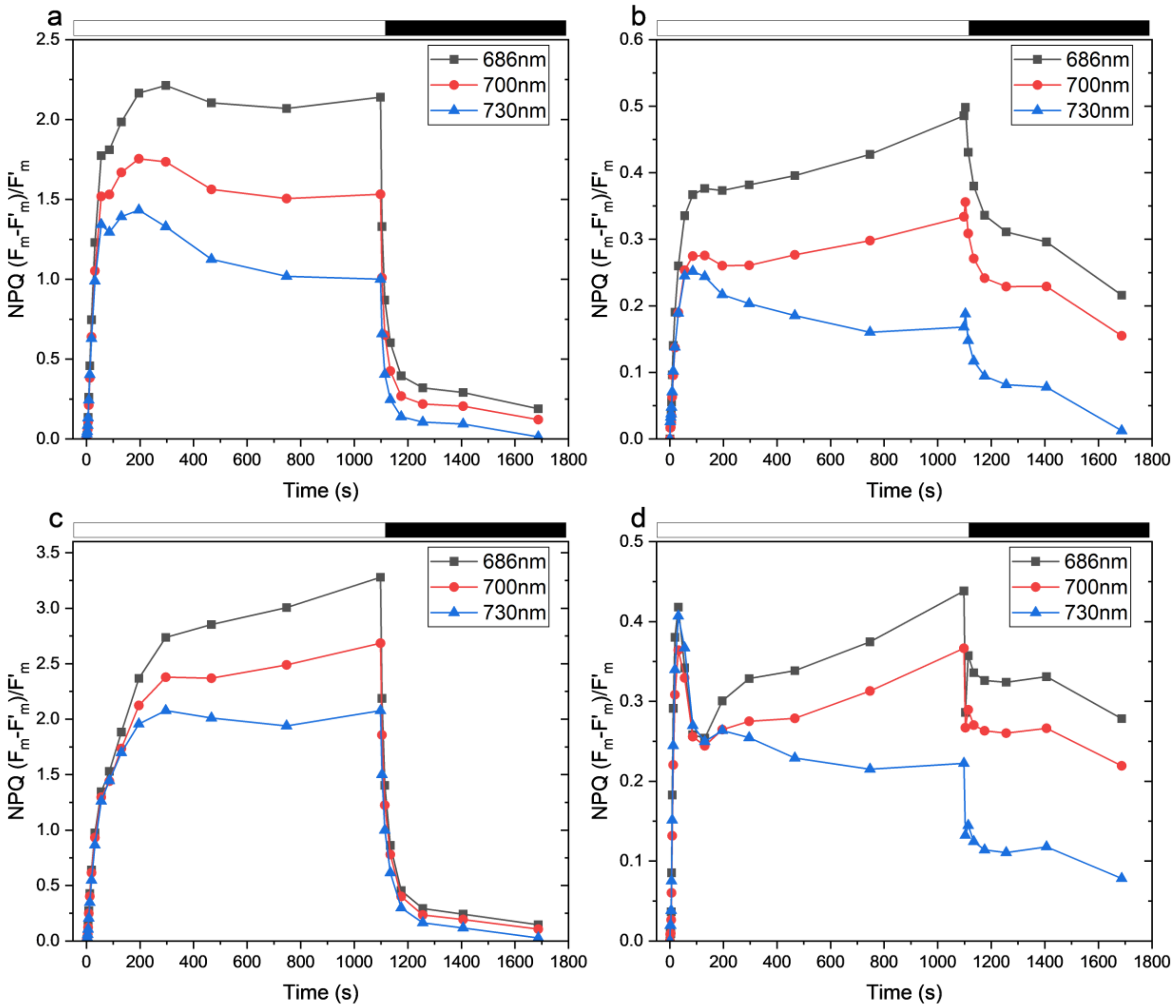


**Fig. S8** NPQ induction and relaxation kinetics of hybrid aspen (**a**) wild type, (**b**) *psbs*, (**c**) oePsbS and (**d**) *vde*. All samples were induced with 600 µmol m$^{-2}$ s$^{-1}$ red actinic light.

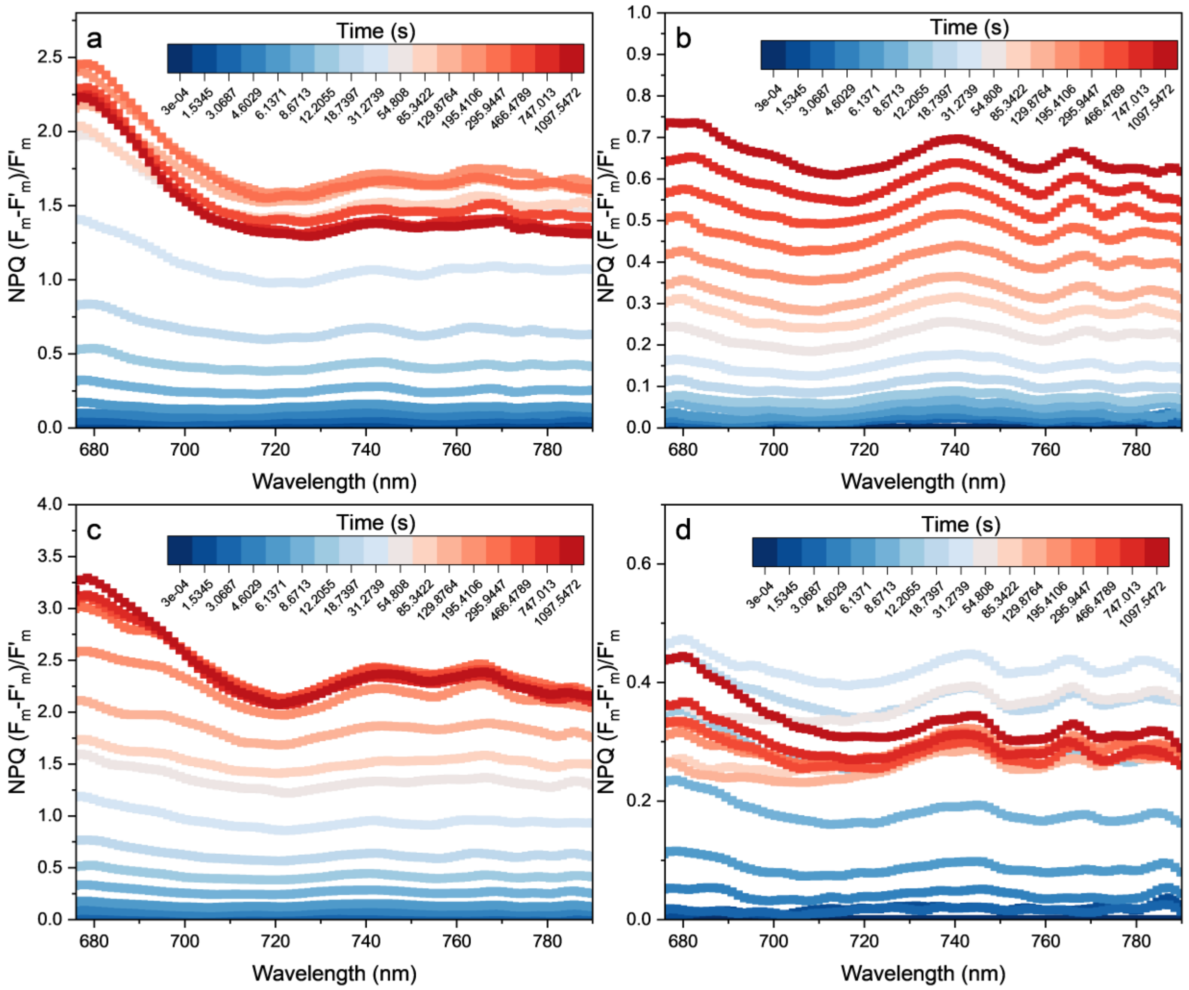


**Fig. S9** NPQ spectra of hybrid aspen (**a**) wild type, (**b**) *psbs*, (**c**) oePsbS and (**d**) *vde*. All samples were induced with 600 µmol $m^{-2}$ $s^{-1}$ red actinic light.

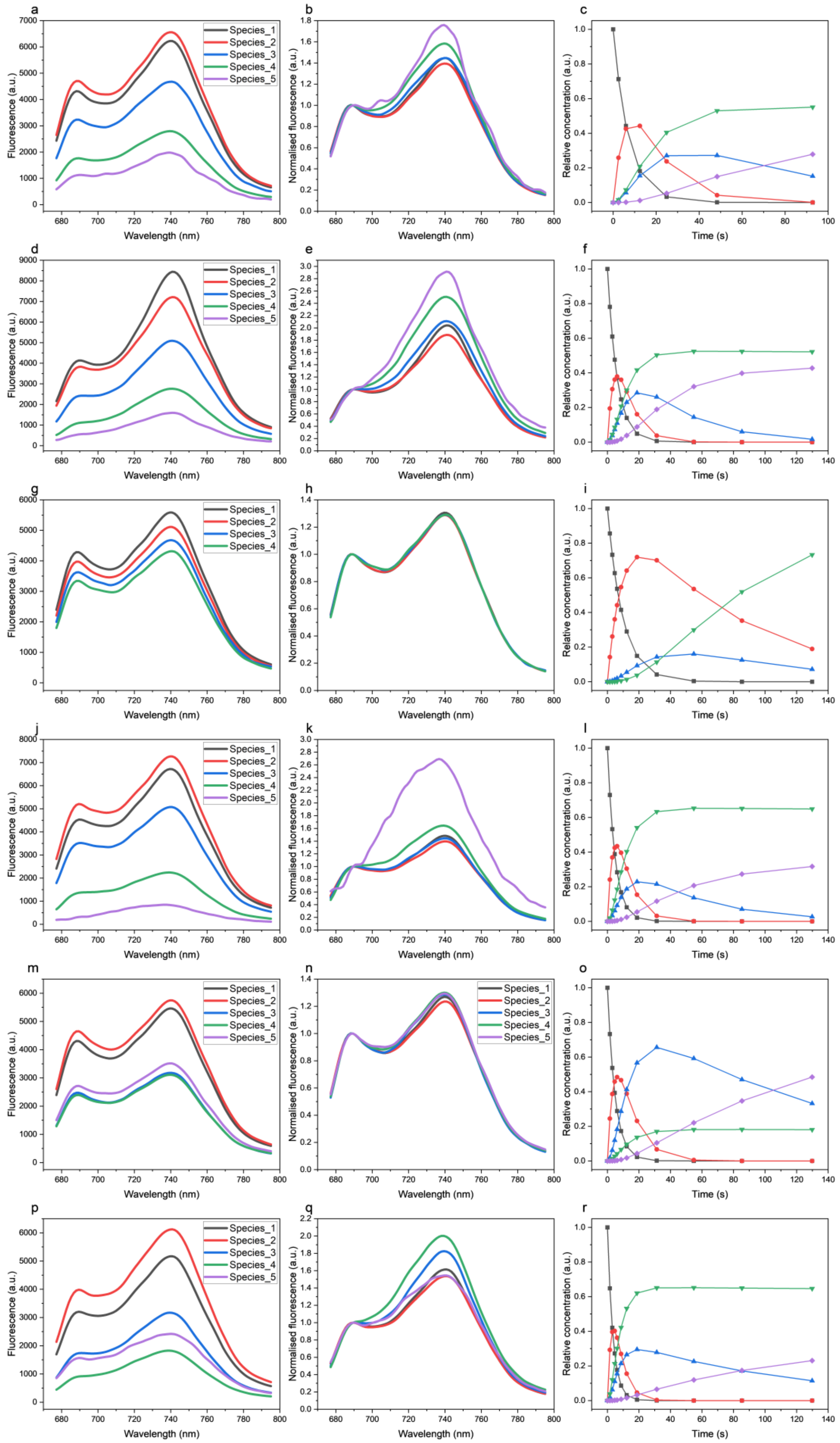


**Fig. S10** Results from global taget analysis of 3D fluorescence spectra for *Arabidopsis* lines as SAES, normalised SAES at 688nm and time dependent concentration profiles respectively in Col-0 (**a**, **b**, **c**,); *P. sylvestris* (**d**, **e**, **f**); *npq4* (**g**, **h**, **i**); L17 (**j**, **k**, **l**); *npq1* (**m**, **n**, **o**) and *npq2* (**p**, **q**, **r**). All samples were induced with 600 µmol $m^{-2}$ $s^{-1}$ red actinic light. The same color code is used for representing the respective species in the various parts of the figure. Species 1 represents the dark-adapted state present at time=0.

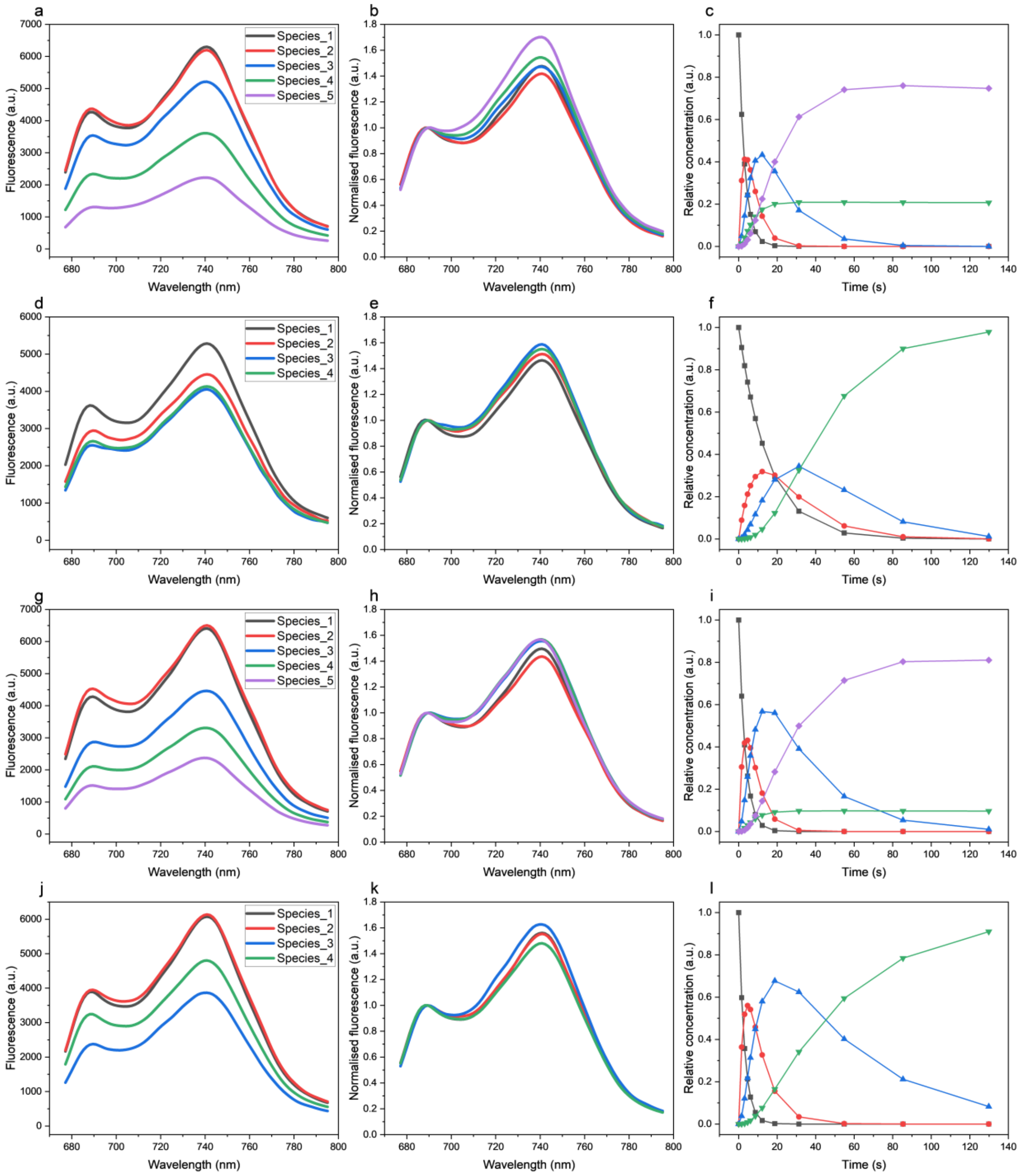


**Fig. S11** Results from global target analysis of 3D fluorescence spectra for hybrid aspen lines as SAES, normalised SAES at 688nm and time dependent concentration profiles respectively in wild type (**a**, **b**, **c**); *psbs* (**d**, **e**, **f**); oePsbS (**g**, **h**, **i**); *vde* (**j**, **k**, **l**). NPQ on all samples was induced by 600 µmol $m^{-2}$ $s^{-1}$ red actinic light. The same colour code is used for representing the respective species in the various parts of the figure. Species 1 represents the dark-adapted state present at time=0.

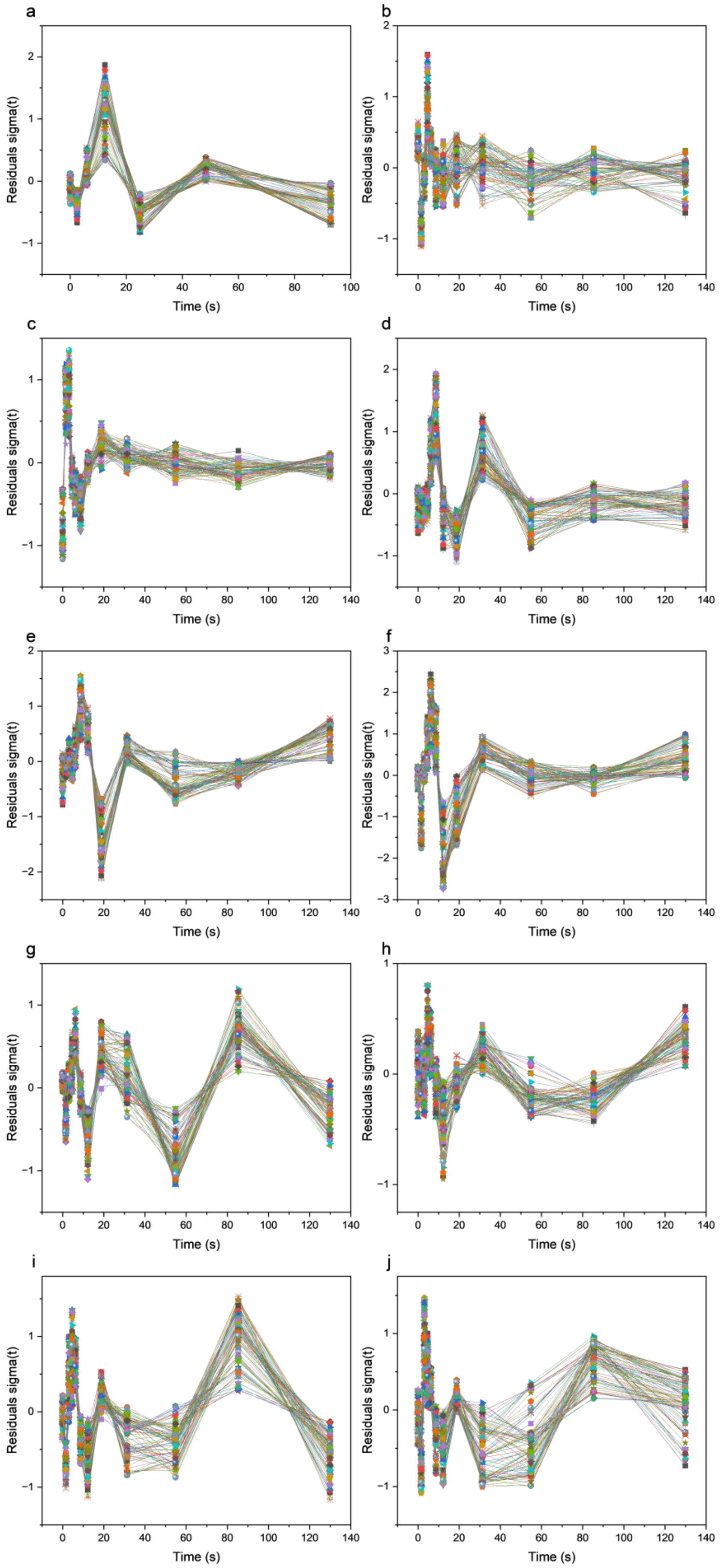


**Fig. S12** Residual plots of global target analysis of *A. thaliana* lines: (**a**) Col-0, (**c**) *npq4*, (**d**) L17, (**e**) *npq1*, (**f**) *npq2*. (**b**) *P. sylvestris*. Hybrid aspen lines: (**g**) wild type, (**h**) *psbs*, (**i**) oePsbS, (**j**) *vde*.

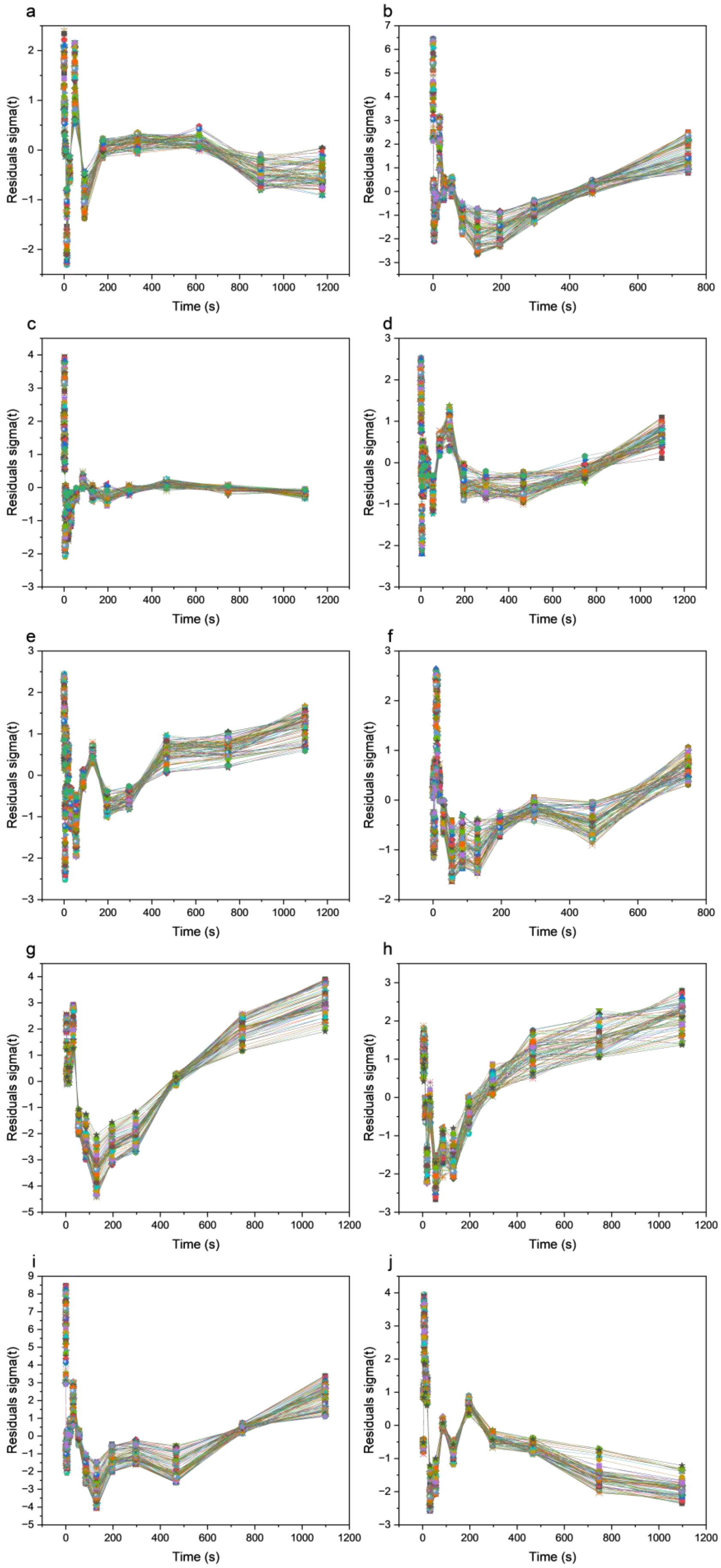


**Fig. S13** Residual plots of MCR-ALS analysis of 3D fluorescence surfaces analyzed over the entire NPQ induction time range. *A.thaliana* lines: (**a**) Col0, (**c**) *npq4*, (**d**) L17, (**e**) *npq1*, (**f**) *npq2*. (**b**) *P. sylvestris*. Hybrid aspen lines: (**g**) wildtype, (**h**) *psbs*, (**i**) oePsbS, (**j**) *vde*. Fits are considered to be good if deviations in the residual plots for the large majority of data points are within the ±2 sigma range. For a few cases (e.g. b, i, and j) the first few data points are substantially exceeding that limit. In these cases for the final fitting of SAES and concentration profiles the first 2 or 3 data points have been discarded.

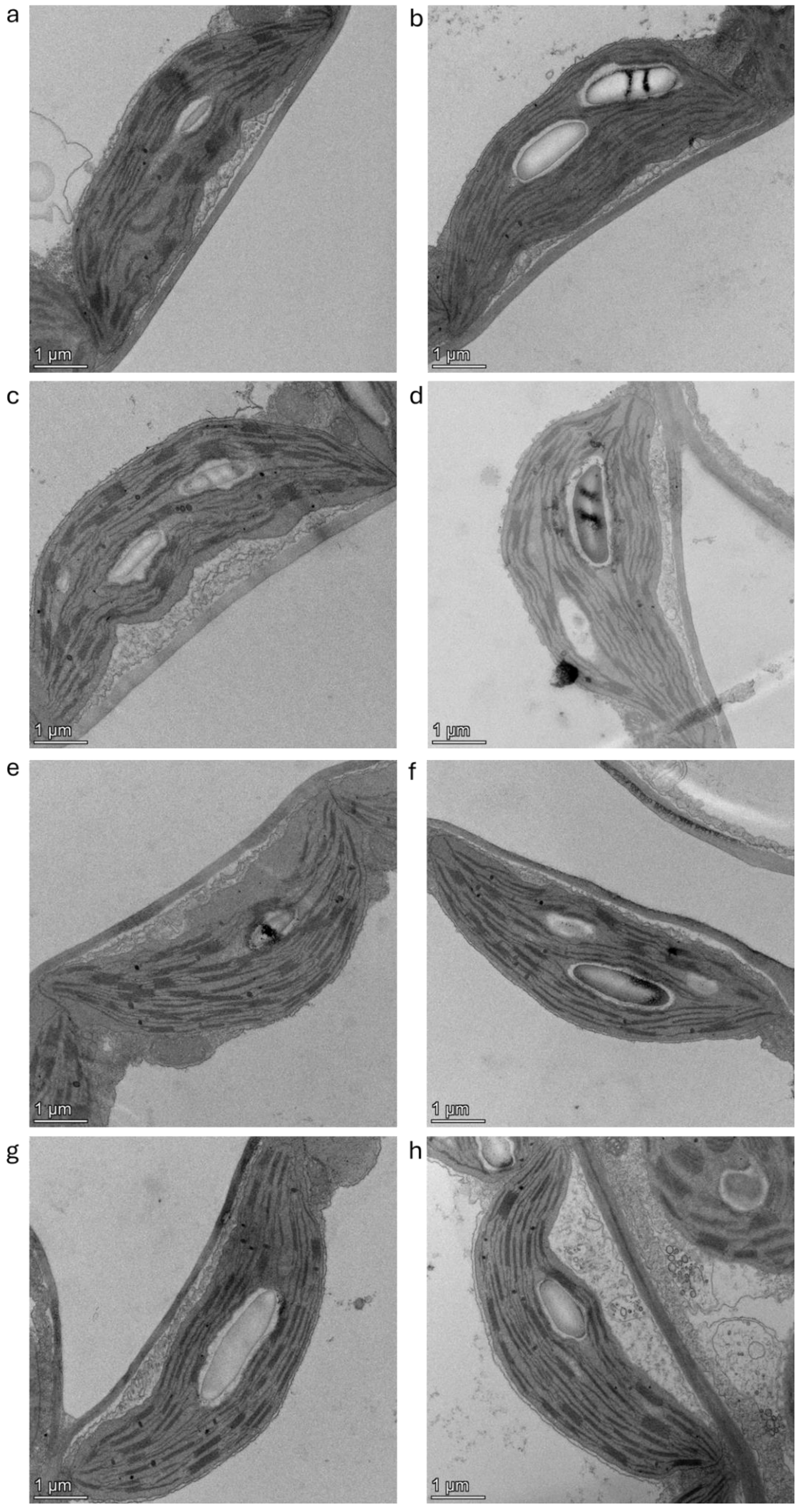


**Fig. S14**
Representative chloroplast images of *Arabidopsis* lines used for thylakoid ultrastructure analyses (c.f. Fig 4 – 5). *npq4*, L17, *npq1* and *npq2* in D (night adapted, dark) in (a) – (d) and AL (actinic light adapted, 600 µmol $m^{-2}$ $s^{-1}$) in (e) – (f) respectively.

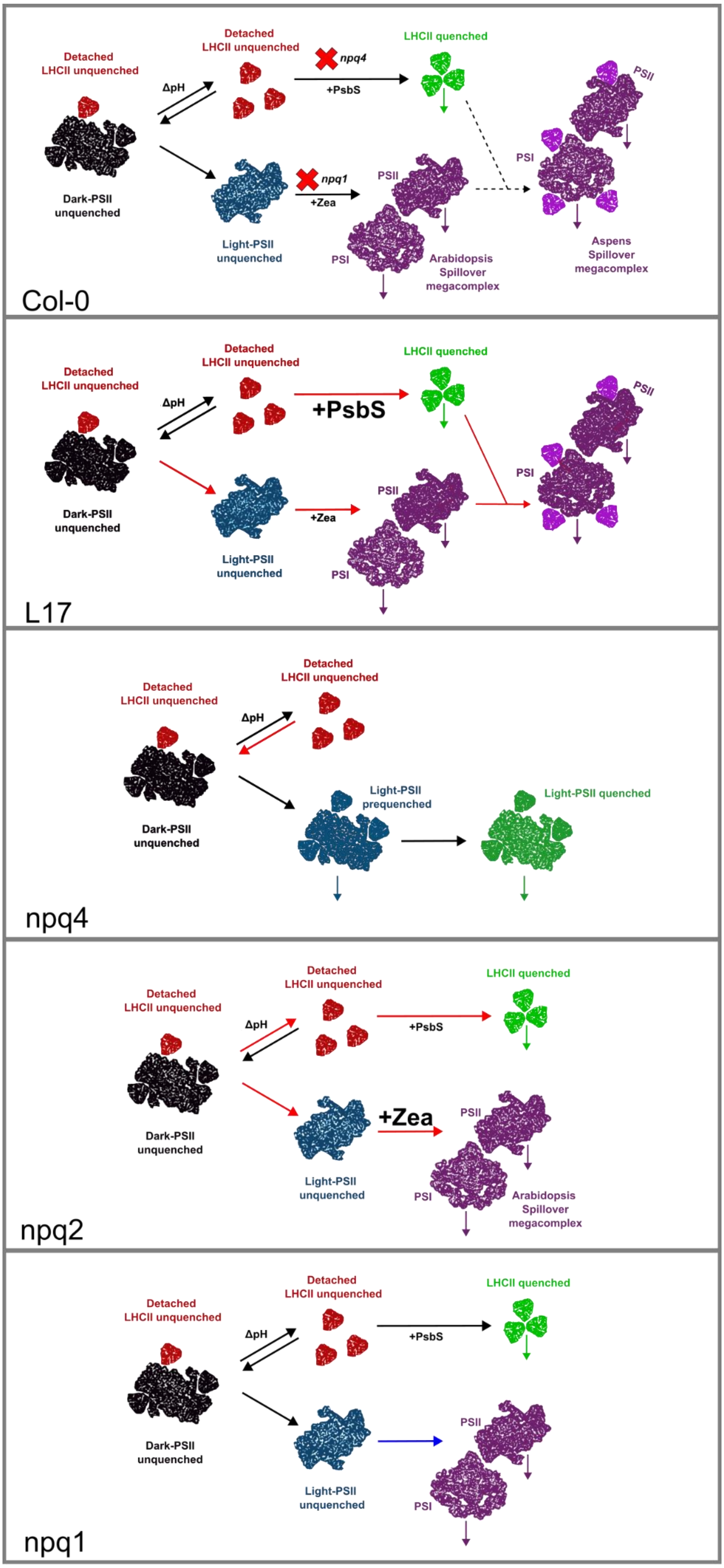


**Fig. S15** Model describing the proposed NPQ mechanism in plants. In Col-0, cooperative work between zeaxanthin and PsbS allows the proper development of quenching species (Fig. 6). When PsbS is overexpressed (L17), the fast development of NPQ (red arrows) leads to the conformation of the quenching complexes present in WT. However, another process seems to take place in L17, where the spillover complex slowly recruits LHCII aggregates, increasing the absorption cross section of the PSII-PSI complexes. The latter resembles the phenomenon that occurs naturally in aspens. When PsbS is absent, the development of new emitting species is affected. In this scenario, PSII changes from an unquenched conformation to a quenched confirmation. In this complex, energy is most likely quenched in LHCII through chl-car and chl-chl charge transfer and charge recombination in their RC. The other master regulator in this process, zeaxanthin, also plays a role in PSII-PSI spillover complex formation. Thus, its relative increase (*npq2*) accelerates the rate of spillover complexes formation (red arrows), whereas its decrease (*npq1*) slows down this process (blue arrows).

| Mutant | sgRNAs sequence | Targeted gene | Targeted gene ID (T89) | Genotyping primers | | | |
|---|---|---|---|---|---|---|---|
| *psbs* | gTCGTTTTAGGGAGACCGTT | *PHOTOSYSTEM II SUBUNIT S (PsbS)* | Potrx000498g00392 | FW | GTCACTCTTGTTGCCTGTCA | RV | GTTTTCCTCGCCCTCTTT |
| | GAACACGATGGCTCAAACCA | | | | | | |
| *vde* | GCAGAGGCTTCATTTCCATG | *VIOLAXANTHIN DE-EPOXIDASE 1 (VDE)* | Potrx034569g11078 | FW | TCTATCTCTCTCTCCCTCTGTG | RV | GGGAAATCCCAAGCACATAGG |
| | GATATTGGCCAGCAGTGCAG | | | | | | |

**Table S1** sgRNA & genotyping primer sequences used for generating and testing CRISPR mutant lines.
